# Internet-delivered cognitive behavioral therapy is associated with distinct blood small RNA profiles in panic disorder

**DOI:** 10.64898/2026.09.03.749122

**Authors:** Shani Vaknine Treidel, Asala Halaj, Lee Mitzva, Tanya Goltser-Dubner, Asher Strauss, Bronislava Gelchinsky-Karagichev, Dalya Pevzner, Osnat Oz, Reaan Amer, Michal Lavon, Ofer Gadish, Estelle R. Bennett, David S. Greenberg, Liran Carmel, Salomon Israel, Ronen Segman, Jonathan D. Huppert, Hermona Soreq

## Abstract

Panic disorder associates with altered fear circuitry, autonomic dysregulation and stress-related immune perturbations, yet its molecular and treatment-response signatures remain poorly defined. Internet-delivered cognitive behavioral therapy (iCBT) is an effective first-line treatment with unexplained outcome variations across individuals. Here, we report altered profiles of small non-coding RNAs in peripheral blood mononuclear cells from panic disorder patients before and after iCBT, compared with longitudinally sampled healthy controls. Thirteen top-ranking microRNAs discriminated patients from controls and separated pre- and post-treatment states, both of which differed from controls. Correspondingly, baseline hsa-miR-125b-5p levels were significantly associated with post-iCBT symptom improvement. Transfer RNA-derived fragments (tRFs) showed even greater discriminatory power, with nuclear-Val-3′-tRF family members exhibiting a pronounced post-treatment shift. Pathway analyses of panic disorder-associated and CBT-responsive small RNA targets converged on immune-related pathways, supporting a role in inflammatory mechanisms and highlighting blood small non-coding RNAs as promising biomarkers of panic disorder and its therapeutic response.

## Introduction

Panic disorder is a disabling anxiety disorder characterized by recurrent unexpected panic attacks and persistent concern about future attacks or their consequences^1^, with lifetime prevalence estimates of 6.2% among women and 3.1% among men in the U.S. population^2^. Although diagnosed clinically, panic disorder is increasingly linked to measurable biological correlates, including altered brain structure and connectivity^3,4^, inflammatory signaling^5,6^, altered DNA methylation^7^, and hormonal and metabolic pathways^2^. Together, these characteristics suggest that molecular parameters may help define vulnerability to panic disorder as well as its symptom state and treatment response.

Cognitive behavioral therapy (CBT), combining psychoeducation, cognitive restructuring and interoceptive and situational exposure, is a first-line treatment for panic disorder^8^. Internet-based CBT (iCBT) delivers these components through structured online modules and has repeatedly shown efficacy^9^, with response or remission rates often around 50% or more^10^ and outcomes comparable to group or face-to-face CBT in several settings^11^. Yet patient response varies, and how molecular changes relate to clinical improvement remains unclear.

Growing evidence suggests that CBT effects may extend beyond symptom improvement to measurable biological changes. In panic disorder, fMRI studies showed CBT’s association with reduced neural threat reactivity and anticipatory-threat responses^12,13^. Poorer CBT response was associated with higher cortisol concentrations, suggesting involvement of the HPA axis^14^. Successful exposure-based CBT was accompanied by partial normalization of MAOA hypomethylation, in contrast to declining methylation levels in non-responders^15^. Other longitudinal studies reported CBT-related changes in DNA methylation, immune-cell composition and serotonin- or immune-related loci^15^, and meta-analyses of anxiety and depression trials have linked CBT to improved immune function^16^. Although these findings do not prove a mechanism of action, they indicate that psychological therapy can leave detectable biological traces.

Small noncoding RNAs are especially relevant in this context because they regulate gene expression, respond to stress and can be measured in blood. MicroRNAs (miRs) are small single-stranded RNAs approximately 22 nucleotides long that bind target mRNAs, thereby reducing transcript stability or translation^17^. Transfer RNA-derived fragments (tRFs), also termed tsRNAs or tDRs, arise from non-random cleavage of precursor or mature tRNAs and can regulate target mRNAs or interact with RNA-binding proteins to regulate translation, stress responses, RNA stability and intercellular signaling^18–20^. Both RNA classes may therefore integrate genetic background, immune state, environmental stress and treatment-related change^21–23^.

MiR dysregulation has been reported in several mental disorders including depression, bipolar disorder, schizophrenia and anxiety-related conditions^24^. In panic disorder, genetic studies have shown associations with miR-tagging single nucleotide polymorphisms involving anxiety-related genes such as BDNF, MAOA, and HTR2C^25^. Direct blood-based studies using serum and plasma identified altered miR levels, including miR-1297, miR-4465, and miR-138-2-3p, implicating GABAergic and glutamatergic pathways and, in some cases, correlating with symptom severity^26,27^. However, data on miRs related to panic disorder treatment remain limited and focus on pharmacotherapy, where sertraline treatment was associated with changes in circulating miRs that correlated with symptom improvement^28^. Whether miRs are altered following CBT or iCBT in panic disorder remained largely unknown.

Compared with miRs, tRFs have only recently been explored in mental disorders such as schizophrenia^29^ and bipolar disorder^30^. In major depressive disorder^31,32^, serum tRF profiles have been proposed as diagnostic biomarkers^31^, and peripheral tRF levels changed following duloxetine treatment in relation to treatment response^32^. Our previous work showed that prenatal stress exposure can affect newborn umbilical cord blood tRF-family patterns^33^. However, tRF levels have not yet been studied in panic disorder or in relation to CBT or iCBT response.

In our current study, we profiled blood miRs and tRFs in patients with panic disorder before and after iCBT, alongside matched controls sampled twice over a similar interval. We asked whether circulating small RNAs can distinguish patients from controls and whether specific miRs or tRFs are associated with iCBT response and symptom improvement. Although no individual miR or tRF passed transcriptome-wide FDR correction, the expression patterns of specific miRs and tRFs distinguished patients from controls and separated pre- from post-iCBT states, with post-treatment remitters displaying a molecular profile distinct from controls. Baseline hsa-miR-125b-5p levels were associated with post-treatment symptom severity, whereas a Nuclear-Val-3′-tRF family signature was associated with long-term symptom severity and differentiated panic disorder and post-iCBT states. Further, target analyses indicated that panic disorder- and iCBT-responsive small RNAs converged on immune and inflammatory pathways, consistent with sex-dependent changes in selected inflammatory transcripts. Together, these findings identify circulating small-RNA signatures as candidate biomarkers of panic disorder and iCBT outcome.

## Results

Thirty-four panic disorder patients participated in a five-module iCBT program and donated blood before treatment (pre-iCBT; n females = 25, n males = 8) and after therapy completion (post-iCBT; n females = 18, n males = 6). Panic Disorder Severity Scale-Independent Evaluator (PDSS-IE)^34^ scores were obtained at intake, post-iCBT, and at three-, six-, and 12-month follow-up interviews. Eighteen controls (n females = 14, n males = 4), matched to panic disorder participants by sex, age, and BMI, donated two blood samples over a similar interval without iCBT intervention (Fig. 1a, b). The groups differed significantly only in baseline PDSS score (Fig. 1c). Peripheral blood mononuclear cells (PBMCs) were isolated and RNA extracted, with sequenced samples chosen only from female panic disorder patients who achieved remission after iCBT (n = 14) and their matched controls (n = 14) at both time points (Supplementary Fig. 1a), reflecting both the higher prevalence of panic disorder in women^2^ and limited male enrollment. After PCA-based outlier removal the final sequencing cohort included panic pre-iCBT (n = 10), panic post-iCBT (n = 11), control pre (n = 12), and control post (n = 10) samples (See Methods, Supplementary Fig. 1b, c).

**Figure 1.**
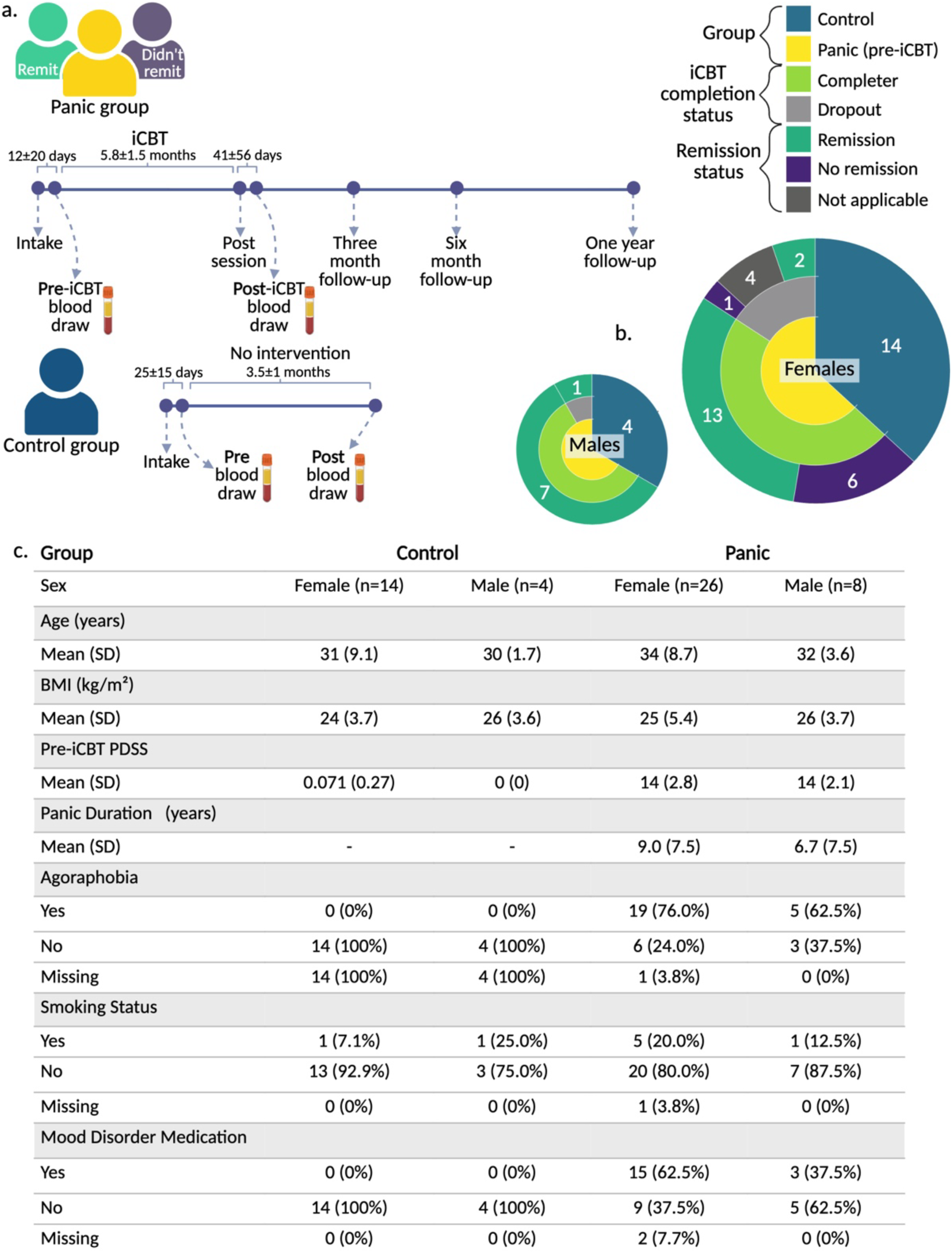
Experimental design. (a) Panic disorder patients and matched controls followed similar sample-collection pipelines; only panic patients underwent internet-based cognitive behavioral therapy (iCBT) and subsequent follow-up interviews. (b) Distribution of female and male participants at the pre-iCBT time point by study group designation, iCBT completion status and remission status (determined at the post-iCBT session). (c) Participant characteristics by group. Only PDSS scores differed significantly between panic and control groups (FDR = 1.1e-24). Created with Biorender.

### Specific miRs differentiate panic disorder and post-iCBT states in females

Sequencing analysis of female participant PBMC samples was carried out as detailed in the Methods (Fig. 2a). Differential expression analysis across the six panic-, treatment-, and time-related comparisons did not identify individual miRs that passed the FDR threshold (Supplementary Fig. 2a-g). We therefore used a ranking and expression-pattern approach to prioritize miRs with consistent directionality across comparisons, while excluding those with marked longitudinal change in controls (see Methods, Supplementary Fig. 2h). This analysis identified 13 miRs: three with a panic disorder-related effect and ten with a transient pattern, defined as opposite changes in panic-related and iCBT-related comparisons (Fig. 2b; Supplementary Fig. 2h). PCA based on these 13 miRs separated panic disorder patients from controls and differentiated pre-from post-iCBT panic samples (Fig. 2c). PC contribution analysis identified hsa-miR-412-5p and hsa-miR-23a-5p as the strongest contributors to patient-control and pre- versus post-iCBT separation, respectively (Supplementary Fig. 3a, b; Supplementary Results). Post-iCBT remitters formed an almost completely distinct cluster rather than showing similarity to controls, suggesting that clinical improvement was accompanied by an altered miR state rather than a return to baseline.

**Fig 2.**
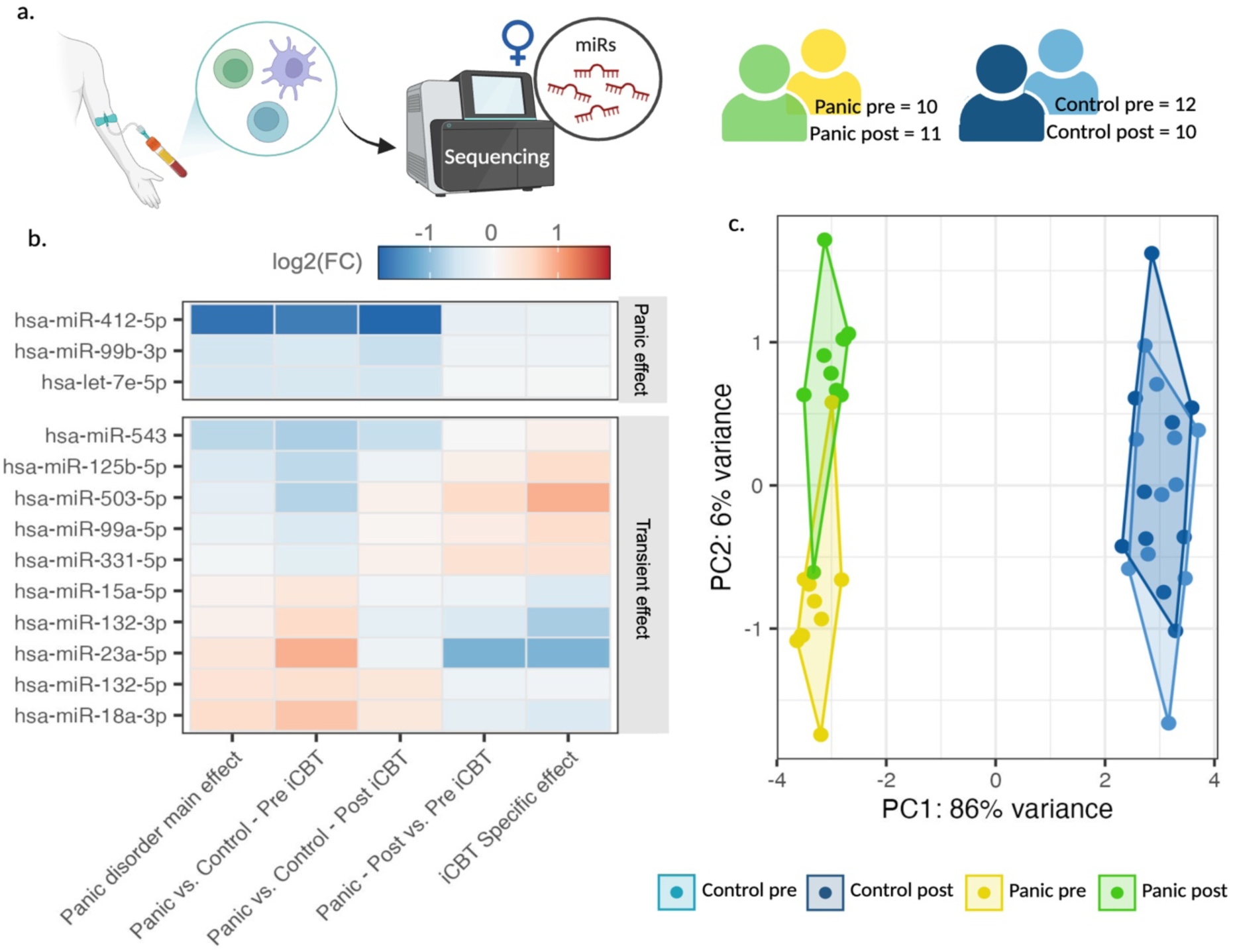
Specific miRs distinguish female controls from panic patients and reflect the response to iCBT. (a) Blood was drawn from participants at two time points, before and after iCBT and RNA was extracted from PBMCs. Female panic patients who achieved remission at the post-iCBT session and their matched controls were sequenced. (b) Heatmap of top-ranked miRs across five differential-expression comparisons, identifying either a panic effect (upper panel) or a transient effect (lower panel) reflecting the difference between control and panic groups before and after iCBT. (c) PCA of the four sequenced groups based on the 13 miRs shown in the heatmap. Created with Biorender.

### Baseline hsa-miR-125b-5p reflects symptom improvement

Based on sequencing expression, effect-pattern classification and contribution to PCA separation, four candidate miRs were selected for qPCR assessment in the full cohort (Supplementary Fig. 3c; Supplementary Results). We then focused on hsa-miR-125b-5p, which was the only one significantly associated with iCBT treatment response by qPCR in the full cohort (Fig. 3a). Female non-remitters had higher baseline hsa-miR-125b-5p levels than female remitters (FDR = 0.022), with a similar post-iCBT trend (FDR = 0.056), whereas male remitters showed lower post-iCBT levels than controls (FDR = 0.042; Fig. 3b). Baseline hsa-miR-125b-5p levels were also associated with post-iCBT PDSS scores after adjustment for age, sex, and BMI (Fig. 3c; Δadjusted R² = 0.36, FDR = 0.026). Related associations showed the same direction for pre-to-post PDSS change (Fig. 3d; Δadjusted R² = 0.25, FDR = 0.054) and 12-month PDSS (Fig. 3e; Δadjusted R² = 0.28, FDR = 0.054). These findings point to baseline hsa-miR-125b-5p as a candidate marker of treatment response and post-treatment symptom severity.

**Fig 3.**
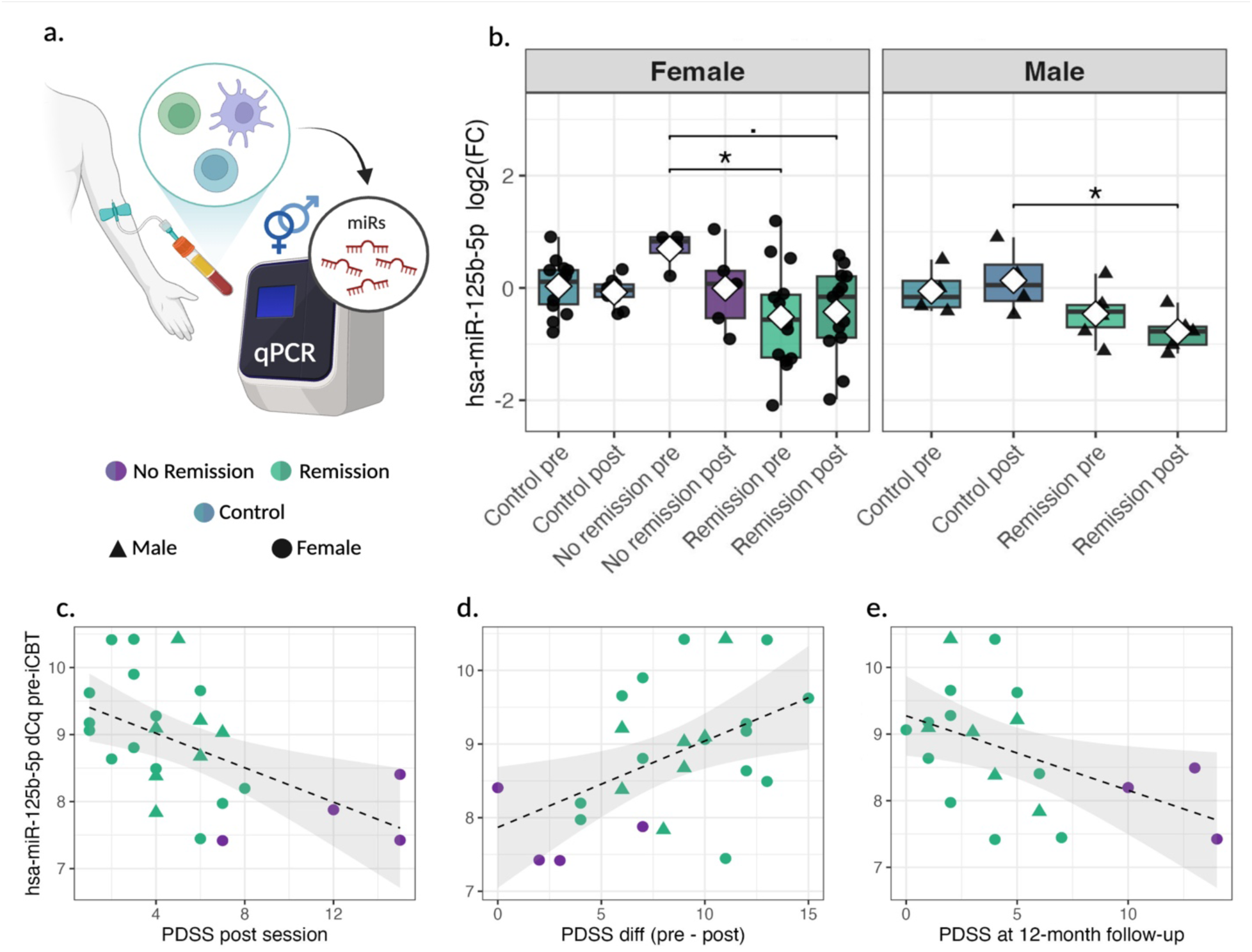
hsa-miR-125b-5p may reflect anxiety improvement following iCBT. (a) PBMC RNA was used to quantify miR levels by qPCR in all male and female participants. (b) Boxplots showing hsa-miR-125b-5p log2FC relative to controls across sex, group, and post-session remission status. Statistics represent Tukey–Kramer tests following ANOVA, with FDR ≤ 0.05 (. = p≤0.1, * = p≤0.05). White diamond in each box indicates mean and the horizontal line marks the median. Sample sizes by group for pre and post time points, respectively: females, remission n = 14/13, no remission n = 4/5, controls n = 14/10; males, remission n = 7/5, no remission n = 0/0, controls n = 4/4. (c–e) Scatter plots showing pre-treatment hsa-miR-125b-5p ΔCq values in relation to (c) PDSS at post-iCBT session, (d) change in PDSS from pre- to post-iCBT session, and (e) PDSS at 12-month follow-up. Points are colored by remission status at the post-iCBT session for (c,d) and at the 12-month follow-up for (e) as in a. Associations were tested using multiple linear regression models with age, sex, and BMI as covariates. Model fit is reported as Δadjusted R² (adjusted R² of the full model minus adjusted R² of the reduced model without ΔCq) and FDR: (c) Δadjusted R² = 0.36, FDR = 0.026; (d) Δadjusted R² = 0.25, FDR = 0.054; (e) Δadjusted R² = 0.28, FDR = 0.054. Dashed lines indicate fitted regression lines and shaded areas indicate 95% confidence intervals. Created with Biorender.

### Specific tRFs separate panic and post-iCBT states in females

Next, we applied a similar analytical framework to tRFs (Fig. 4a). As in the miR analysis, differential expression analysis did not identify individual tRFs that passed the FDR threshold (Supplementary Fig. 4a-g). However, the ranking and pattern-based workflow identified 13 altered tRFs, including one panic-effect and 12 transient-effect tRFs (Fig. 4b; Supplementary Fig. 4h). PCA based on these tRFs was similar to that of the miRs but showed stronger group separation, again placing post-iCBT remitters in a distinct cluster from controls (Fig. 4c) and supporting a distinct post-iCBT molecular state. PC contribution analysis identified mtDR-35:66-His-GTG-1, tDR-1:32-Lys-TTT-3-M2, and tDR-L1:34-His-GTG-1 as the leading contributors to group separation (Supplementary Fig. 5a-b), although qPCR assessment in the full cohort did not show significant differences (Supplementary Fig. 5c), possibly because closely related tRFs often differ by only a few nucleotides, making qPCR discrimination difficult^35^. These results suggest that specific tRFs capture panic disorder- and treatment-associated changes that are not detected by individual differential expression analysis but become informative as coordinated group-level patterns.

**Fig 4.**
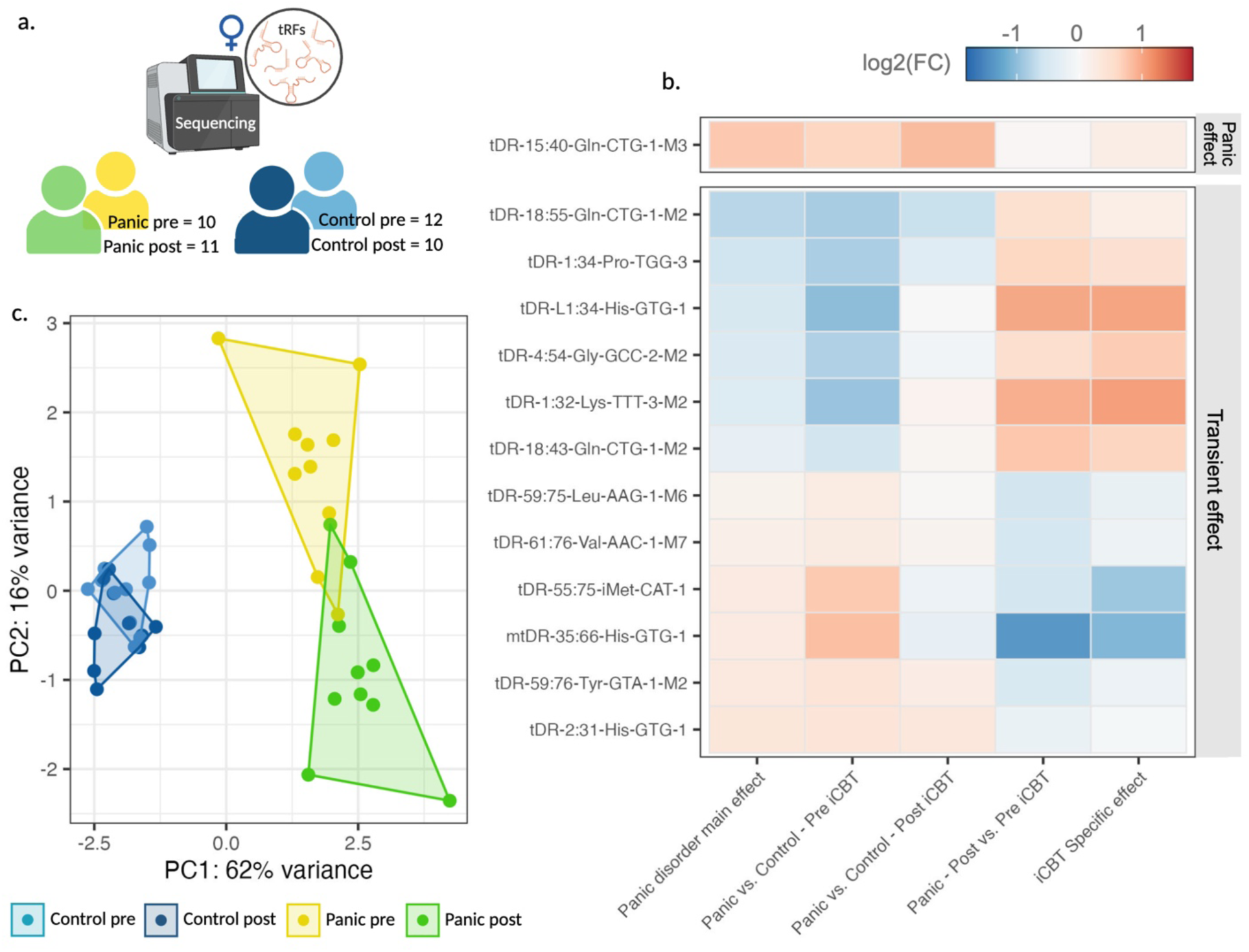
Specific tRFs distinguish controls from panic patients and reflect the response to iCBT. (a) tRF sequencing results are shown for female panic patients who achieved remission at the post-iCBT session, and their matched controls. (b) Heatmap of top-ranked tRFs across differential expression comparisons, identifying either a panic effect (upper panel) or a transient effect (lower panel) between control and panic groups before and after iCBT. (c) PCA of all sequenced participants based on the 13 tRFs shown in the heatmap. Created with Biorender.

### tRF-family analysis highlights Nuclear-Val-3′-tRFs

Since tRFs may act in coordinated families, as observed in our prenatal-stress study^33^, we next grouped individual tRFs by genomic origin, parental tRNA-encoded amino acid and cleavage type (Fig. 5a). Exact binomial testing for family-consistent directionality identified 16 altered tRF families across the six differential expression comparisons (FDR ≤ 0.05; Supplementary Fig. 6). After excluding families that changed significantly in the longitudinal control comparison, two panic-related nuclear tRF families remained: Val-3’-tRF and His-i-tRF (Fig. 5b).

**Fig 5.**
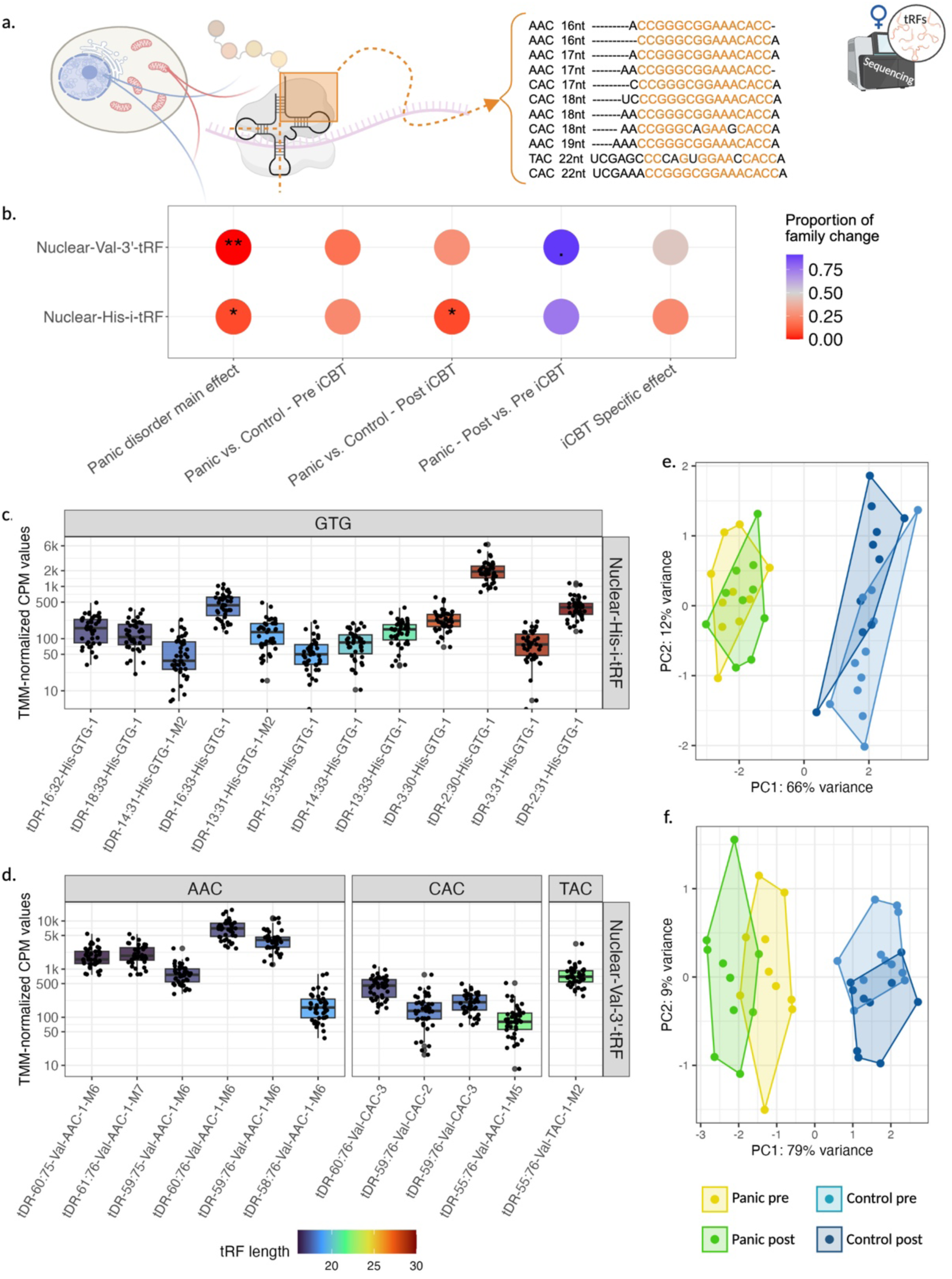
Specific tRF families distinguish pre- and post-iCBT panic groups from controls. (a) Illustration of tRF family grouping, shown for the Nuclear-Val-3′-tRF family. tRFs can be grouped into families that share sequence motifs, defined by genomic origin, parental tRNA amino acid and cleavage type. (b) Dot plot of two tRF families showing directional family-level changes assessed with exact binomial tests. Dot size indicates the number of individual tRFs within each family and dot color indicates the proportion of family members increased or decreased in each differential expression comparison. Only families with FDR ≤ 0.05 in at least one of the five differential expression analyses and non-significant results in the control post- vs. pre-comparison are shown. (c) Boxplot showing expression levels of individual tRFs in the Nuclear-His-i-tRF family highlighted in (b). Each point represents a participant; color indicates tRF length (ranging from X to 30 nucleotides), with the parental tRNA codon in the grey area above the plots. Data are plotted on a logarithmic scale with major ticks at intervals of 1, 2 and 5. (d) Boxplot as in (c) showing Nuclear-Val-3’-tRF family. (e) PCA plots based on Nuclear-His-i-tRF family members. (f) PCA based on Nuclear-Val-3′-tRF family members. Sample sizes: panic pre-iCBT n = 10, panic post-iCBT n = 11, control pre n = 12, control post n = 10. Created with Biorender.

The Nuc-His-i-tRF family contained variable-length fragments, consistent with broader i-tRF heterogeneity^36,37^, and differentiated panic and control groups but did not separate pre- from post-iCBT panic samples (Fig. 5c, e). By contrast, the Nuc-Val-3’-tRF family, comprising 11 short fragments from three valine tRNAs, showed a coherent length range and shared a sequence motif (Fig. 5a, d). One family member, tDR-61:76-Val-AAC-1-M7, also belonged to the transient-effect tRF group (Fig. 4b), increased in panic patients relative to controls and decreased after iCBT. PCA based on these Nuc-Val-3’-tRFs separated pre- and post-iCBT panic groups from controls. Here as well, along PC1, post-iCBT remitters occupied a distinct molecular position rather than shifting toward controls (Fig. 5f). This family therefore emerged as a potential tRF signature of panic disorder and the post-iCBT state, further supporting the notion that remission may be accompanied by a distinct molecular state rather than a return to control levels.

### Post-iCBT Nuc-Val-3′-tRFs associate with longer-term anxiety

To further investigate the Nuc-Val-3′-tRF family we used qPCR to measure the levels of two family members, tDR-61-76-Val-AAC-1-M7 and tDR-59-76-Val-AAC-1-M6, in the full cohort. These two tRFs were selected as short, motif-containing representatives of their family, with one derived from the transient-effect set and the other chosen as a less biased, highly expressed family member. For both tRFs, similar associations were seen between post-iCBT levels and later PDSS scores (Supplementary Fig. 7a-h), consistent with the idea that these assays capture a shared Nuclear-Val-3′-tRF family signal rather than fully independent fragments. Associations were strongest at six- and 12-month follow-up interviews: tDR-59:76-Val levels were significantly altered at six months (Δadjusted R² = 0.27, FDR = 0.030), and both tDR-61-76-Val and tDR-59:76-Val showed a significant association at 12 months (tDR-61:76-Val: Δadjusted R² = 0.42, FDR = 0.038; tDR-59:76-Val: Δadjusted R² = 0.44, FDR = 0.030; Supplementary Fig. 8d, g, h). Both showed weaker but directionally consistent associations at the post-iCBT and three-month assessments. Overall, lower post-iCBT Nuclear-Val-3′-tRF levels were associated with higher later anxiety, suggesting that this family may reflect maintenance of remission or vulnerability to symptom recurrence.

### Panic- and iCBT-responsive small RNAs converge on immune and inflammatory pathways

To understand the biological consequences of panic disorder- and iCBT-responsive small RNAs, we examined whether they target shared biological pathways. Target enrichment analysis of the 13 panic- and transient-effect miRs (Fig. 2b), based on 2,481 unique experimentally validated targets, revealed 209 significant WikiPathways^38^, 31 of them immune-related, consistent with inflammatory alterations previously reported in panic disorder^5,6^ (Supplementary Fig. 8, Supplementary Results). Since some tRFs can function through a miR-like mechanism^20,35^, we identified Nuc-Val-3′-tRF family targets using a tRF-specific validated target tool^39,40^, identifying 458 genes and 66 WikiPathways^38^. Combining the miR and tRF-family enrichment analyses revealed 40 shared pathways (Supplementary Table 1). Focusing on immune regulation, we constructed a bipartite network linking immune-related WikiPathways to these miRs and tRFs (Fig. 6). This revealed 36 pathways connected by 56 directed interactions between the miRs and the Nuc-Val-3’-tRF family, indicating that independent panic- and iCBT-responsive small-RNA classes can converge on a common immune-regulatory axis.

**Fig 6.**
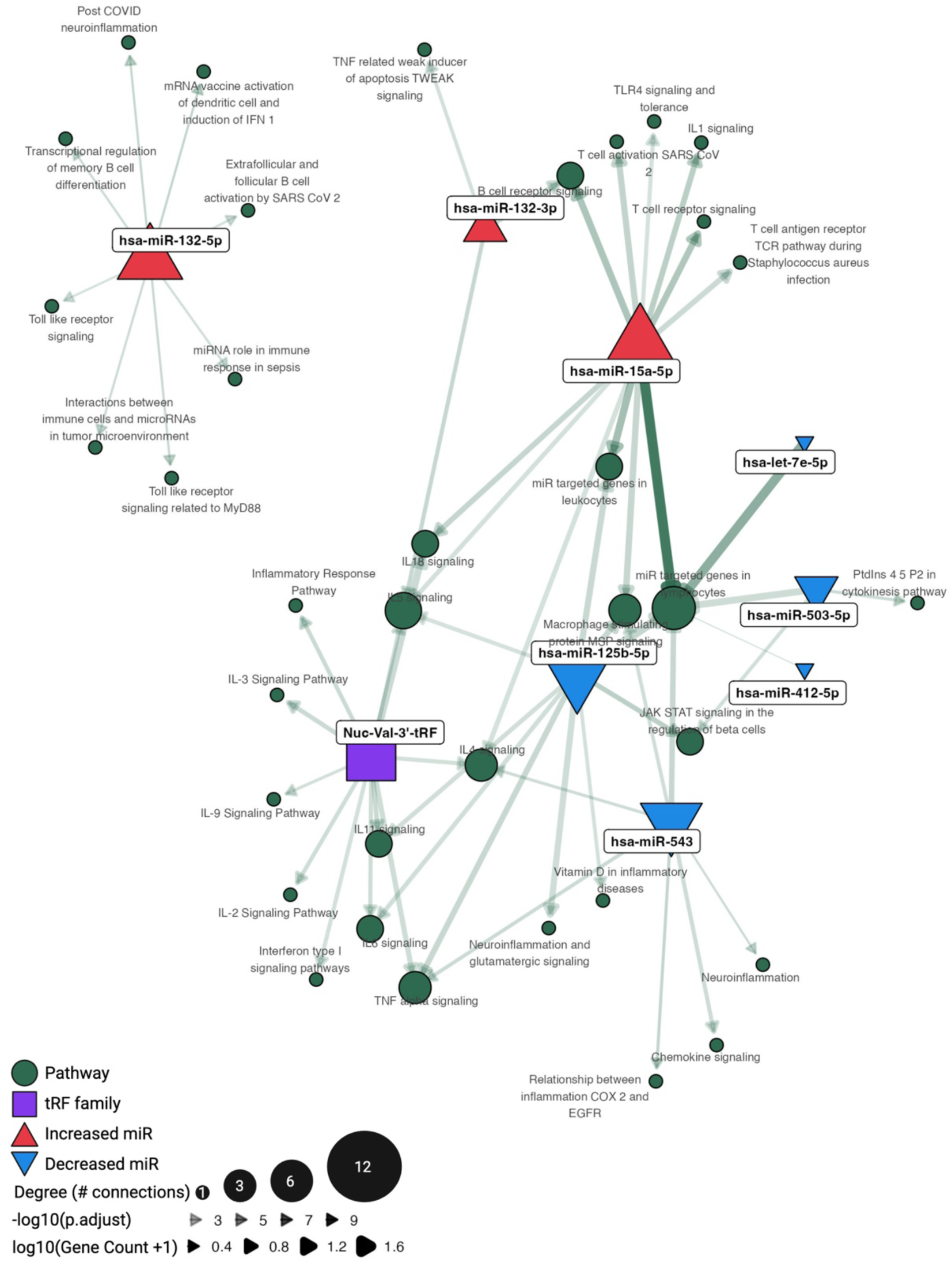
Panic disorder- and CBT-responsive small RNAs regulate immune and inflammatory - related pathways. Bipartite network diagram illustrating the regulatory connections between panic- and transient-effect miRs and the Nuc-Val-3’-tRF family and their enriched immune- and inflammation-related WikiPathways. Gene targets were identified using experimentally validated target databases (miRTarBase for miRs, tRFtar for tRFs), and immune- and inflammation-associated pathways were selected from significant WikiPathways enrichment results for each small RNAs. Pathways are represented by green circles and are sized by degree of connectivity with the small RNAs (based on the number of total connections). Green arrowhead sizes and transparency indicate enrichment significance by −log10(adjusted p-value). Node shapes and colors represent small RNA type and direction of change: upward-pointing red triangles indicate miRs increased in panic pre-iCBT compared with controls, downward-pointing solid blue triangles indicate miRs decreased in the same comparison, and the purple square indicates the Nuclear-Val-3′-tRF family, which was also increased in this comparison. Two miRs (hsa-miR-99b-3p and hsa-miR-99a-5p) that did not target these pathways were excluded from the network. Created with Biorender.

Within this network, increased small RNAs, including hsa-miR-15a-5p, hsa-miR-132-5p, and the Nuc-Val-3′-tRF family, and decreased miRs, including hsa-miR-125b-5p and hsa-miR-543, mapped to overlapping inflammatory, lymphocyte and cytokine-related pathways (Fig. 6). Thus, although specific small RNAs changed in different directions, their predicted targets converged on related immune processes (Supplementary Fig. 8, Supplementary Results). To test this experimentally, we measured seven immune-related mRNA candidates by qPCR in the full cohort. Treatment-associated mRNA changes were sex-dependent, with the clearest signal in male remitters, who showed post-iCBT increases in STAT1 (Tukey–Kramer tests, p.adjusted = 0.043) and IL6R (p.adjusted = 0.035) and lower baseline TGF-beta expression (p.adjusted = 0.023) that reached control levels after treatment (Supplementary Fig. 9). In contrast, female participants showed few changes overall, with the only statistically significant finding being lower baseline IL1b in non-remitters than in healthy controls (p.adjusted = 0.044). These findings are consistent with the pathway analysis and suggest that immune-related transcriptional responses to iCBT may be sex-dependent.

## Discussion

To study small-RNA involvement in panic disorder and iCBT response, we profiled PBMC small noncoding RNAs in panic disorder patients before and after iCBT and in longitudinally sampled matched controls. Top-ranked miRs and tRFs distinguished patients from controls and separated pre- from post-iCBT states. Specifically, hsa-miR-125b-5p emerged as the strongest clinically-linked miR candidate, with baseline levels associated with remission status and post-treatment PDSS outcomes. Among tRFs, the Nuc-Val-3’-tRF family formed a coherent signature that differentiated panic disorder and post-iCBT states and was related to longer-term symptom severity. Together with immune-related target pathway analysis and mRNA measurements, these findings suggest that peripheral small RNAs may reflect both panic disorder- and treatment-linked molecular changes.

A central observation was that clinical remission did not correspond to molecular ‘normalization’. In both miR- and tRF-based analyses, rather than overlapping with controls, post-iCBT remitters formed a distinct group. Although this contrasts with reports of partial normalization in neural measures^12^ and MAOA gene methylation^41^ after CBT, responses may differ across biological processes. Panic disorder has been associated with elevated pro-inflammatory cytokines^5^, and psychosocial interventions can reduce inflammatory markers^16^, yet few studies have examined whether treatment restores immune profiles to control levels. One such study found persistent immune changes in CBT-treated subjects despite clinical remission following CBT and benzodiazepine treatment^42^. These findings, together with the miR- and tRF-targeted immune pathways observed here, may explain why post-iCBT samples formed a separate cluster and suggest that circulating small RNAs may reflect residual immune vulnerability, compensatory regulation and/or persistent molecular traces of prior panic disorder symptoms.

hsa-miR-125b-5p exhibited treatment-response heterogeneity within the panic disorder group, with remitters and non-remitters showing opposite baseline patterns, arguing against a simple disease-state interpretation. MiR dysregulation has been implicated in stress-related psychiatric disorders^24^, and validated hsa-miR-125b-5p targets in our analysis were enriched in immune- and stress-related pathways, including IL6, TNF-alpha, MAPK, JAK-STAT and BDNF-related signaling^2,5,6,43^. The IL6 pathway is particularly relevant because it is implicated in panic disorder^5^ and stress-challenge responses^6^. Our results support hsa-miR-125b-5p as a candidate biomarker for iCBT response, but replication in larger independent cohorts is required.

This study extends panic disorder small RNA research to tRFs. Although individual tRFs did not reach transcriptome-wide significance, the top differentially expressed tRFs separated the study groups, and family-level analysis highlighted Nuc-Val-3′-tRFs as the strongest markers of iCBT response. This further supports the concept that tRFs act as coordinated fragment families rather than individual entities and is consistent with our previous findings^33^. Our results also align with emerging evidence that tRFs play a role in psychiatric disorders including schizophrenia^29^, bipolar disorder^30^ and major depressive disorder^31,32^, as well as neurological conditions^21^.

Several limitations to our study should be considered. First, the sequenced cohort was modest and restricted to female remitters, increasing clinical focus but limiting generalizability. Although qPCR validation included the broader cohort, the male subgroup was small and all male participants achieved remission, preventing assessment of sex-specific non-response. The female non-responder group was also small, further limiting interpretation. Second, PBMC profiles reflect mixed immune-cell populations, so differences may reflect cell-type composition, small-RNA expression within cell types, or both. Third, no individual miR or tRF reached FDR significance in sequencing, and candidate selection relied on ranked and pattern-based approaches. This may reflect the limited sample size, but also a broader challenge in psychiatric research, where biologically meaningful effects are often modest, heterogeneous and difficult to detect with standard measures. Finally, medication was permitted among patients but not controls. Patients’ medication was required to remain stable, making it unlikely to drive small-RNA changes, although it may have affected patient-control comparisons. Controls were sampled longitudinally without an alternative psychological intervention, limiting isolation of iCBT-specific effects from other time- or participation-related effects. Nevertheless, this no-intervention control group enables both separation of treatment-associated changes from temporal variation and the evaluation of whether remission produces a control-like or distinct molecular state. Given the established efficacy of iCBT^10,44,45^, assigning healthy controls to an unnecessary psychological intervention would provide limited interpretive benefit while raising ethical and practical concerns.

In conclusion, this study suggests that panic disorder and iCBT response are accompanied by distinct changes in blood small-RNA profiles. Specifically, hsa-miR-125b-5p emerged as a candidate biomarker of treatment-related heterogeneity and longer-term symptom outcomes, while Nuclear-Val-3’-tRFs revealed a coherent tRF-family signature associated with panic disorder and the post-iCBT state. The convergence of panic disorder- and CBT-responsive small RNAs on immune-related pathways, together with sex-dependent changes in inflammatory mRNAs, supports the involvement of peripheral neuroimmune regulatory networks in panic disorder biology and treatment responses. Larger longitudinal studies integrating small RNAs with immune-cell composition, target mRNA levels, cytokine levels and broader clinical outcomes will be needed to determine whether these biomarkers identify biological vulnerability persisting after clinical remission or, more importantly, predict individual CBT response.

## Acknowledgments

The authors thank all participants for their time, effort, and invaluable contribution to this study. This work was supported by the Prof. Milton Rosenbaum Endowment Fund for Research in Psychiatric Sciences (0341802) and the Hebrew University of Jerusalem Personalized Medicine Fund (036.1575).

## Ethics approval

All participants provided informed consent during the intake interview, under approval from the Hebrew University ethics board (titled “Biological markers in panic disorder and in response to internet-based and face-to-face cognitive behavioral therapy” for panic disorder patients and “Biological markers and emotional aspects in healthy subjects without anxiety disorders” for healthy controls). Additional clinical approval was obtained from the Hadassah Medical Organization Institutional Review Board (IRB) committee, and participants provided a second informed consent form on the day of the blood-draw (0577-18-HMO).

## Data availability

The sequencing data underlying this study will be deposited in the Gene Expression Omnibus (GEO) upon manuscript publication.

## Additional Information

Supplementary figures and tables are available in a separate file.

## Author contributions

S.V.T. recruited participants, collected blood samples, conducted experiments, managed and performed data analyses, and drafted the manuscript. A.H. coordinated participant recruitment, conducted clinical interviews, and supervised the iCBT intervention. R.S. served as the study physician and contributed to study management. T.G.D. coordinated sample processing and contributed to study management. A.S. developed the iCBT treatment used in the study and was a study therapist. B.G.K. contributed to blood collection and participant consent procedures. D.P., O.O., R.A. and M.L. contributed to sample processing. L.M. and O.G. contributed to experimental work and data analysis. E.R.B. contributed to experimental design and execution and participated in manuscript writing and editing. D.S.G. contributed to manuscript writing and editing. S.I. contributed to study management. L.C. provided guidance on statistical and data analyses and contributed to manuscript writing and editing. J.D.H. coordinated all iCBT-related procedures, supervised the iCBT intervention, and contributed to management. H.S. supervised the project and led manuscript writing and editing.

## Methods

### Participant recruitment

Panic disorder patients and control participants were recruited between March 2019 and February 2023 through online and public advertisements and referrals. Advertisement respondents were directed to register through a study website, where they signed an online informed consent form and completed a series of online questionnaires similar to Halaj et al. (2023). Participants were then screened by phone and, if eligible, invited for a clinical intake interview in person or via Zoom. During the intake interview, participants provided informed consent again and were assessed using the Mini International Neuropsychiatric Interview for DSM-5 Axis I disorders ^46^ and the PDSS-IE ^34^. Eligibility criteria for the panic disorder group included a principal DSM-5 diagnosis of panic disorder and/or agoraphobia, age of 18-65 years, panic disorder duration of at least three months, stable panic disorder medication dosage for at least three months before treatment with no dose increase during treatment, access to the internet and willingness to use it.

Control participants were recruited to match panic disorder patients by sex, age (within a six-year range) and BMI (within a four-point range). They underwent a similar pipeline, including online questionnaires followed by an in-person or Zoom clinical intake interview, and were eligible if they did not meet criteria for panic disorder or any other psychiatric disorder. Exclusion criteria for both groups were substance abuse or dependence within the previous six months, active suicide potential within the previous six months, any current or past psychosis or bipolar I disorder, current weekly or biweekly psychotherapy, history of a complete course of panic-focused CBT, neurological disorders with cognitive decline, chronic physical illnesses, or current infectious disease (Fig. 1).

### Internet-delivered CBT (iCBT)

Following intake, panic disorder participants received internet-delivered CBT (iCBT), based on Barlow’s updated treatment protocol^47^. The treatment included six modules and involved a 16-week program. Completion was achieved by completing five or more modules. Participants were supervised by therapists who were MSc- or PhD-level students in the Clinical Psychology program at the Psychology department of the Hebrew University. Therapists were trained in CBT for panic disorder and met weekly with a licensed clinical psychologist for group supervision (JDH). For more details regarding the iCBT protocol, see Strauss et al. (2022). The primary outcome measure was the PDSS-IE^34^, administered by an evaluator blinded to treatment progress. This assessment was conducted at pre- and post-treatment interviews (intake interview and post session, respectively) and at three follow-up interviews (three, six, and twelve months). The cutoff for participating in the study was a PDSS-IE score of ten or above for the panic group, and one or less for the controls. Remission status was assigned to patients with panic disorder at the post-session and follow-up time points if their PDSS-IE score at that time was lower than or equal to seven, with an additional requirement for remission assignment at the post-session: a difference of at least four points between the pre-iCBT and post-iCBT evaluations. Control participants received no intervention between the two blood draws.

### Sample collection

Participants donated blood at two time points: the first after the intake interview (pre-iCBT) and the second after the post-iCBT session, when therapy was concluded, for the panic group, or approximately three months with no intervention after the intake interview for the control group (post-iCBT). Before the COVID-19 pandemic, blood was drawn on the same day as the respective meeting. Once the pandemic started, all interviews became virtual and blood draws were scheduled separately (Fig. 1a). All participants received financial compensation equivalent to $33 for each blood draw and, once the pandemic started, were required to sign a health declaration confirming no COVID-19-related symptoms at the time of blood draw. Venous blood was collected by a medical specialist at the Hadassah Medical Center, Mount Scopus, Jerusalem, into BD Vacutainer ACD Solution A 8.5 ml tubes (#VACU366645). Blood draw was timed to either 10:00 or 16:00 to avoid interference from circadian cortisol peaks. Following collection, blood samples were transported at room temperature to Hadassah Ein Kerem Medical Center, where PBMC separation was performed within two hours.

### PBMCs and RNA extraction

Citrate blood samples were centrifuged with a Histopaque gradient (Sigma-Aldrich Histopaque-1077, H8889) to separate peripheral blood mononuclear cells (PBMCs), which were selected as the preferred blood fraction for the study due to the inflammation-related effects in panic disorder^5,6^. PBMCs were stored in 1 mL of 10% DMSO at -80°C and moved to liquid nitrogen for long-term storage, as previously described^48^. PBMC samples were shipped on dry ice to the Soreq laboratory in the Department of Biological Chemistry, the Institute of Life Sciences, Edmond J. Safa Campus of the Hebrew University of Jerusalem, where they were again kept at -80 °C for short-term storage until RNA extraction. For that process, PBMC samples were rapidly thawed by hand and added to 10 mL RPMI-1640 (Sigma-Aldrich, R0883) preheated to 37°C. Samples were then centrifuged for 10 minutes at 300g at room temperature and the supernatant discarded. RNA was extracted from the pellet immediately, using the miRNeasy Mini Kit (Qiagen, 217004) according to the manufacturer’s instructions. RNA yield was determined (NanoDrop 2000, Thermo Fisher Scientific), with average yields of 317.6 ng/µL, and quality was assessed by standard gel electrophoresis.

### Sequencing and alignment

Only samples from female panic patients who achieved remission post-iCBT and had matched female controls were considered for sequencing (Panic n = 14, Control n = 14), due to the low number of male participants in the study (Panic n = 8, Control n = 4, out of 13 in total), in line with the higher female prevalence of panic disorder^2^. Final samples were selected based on RNA integrity number (RIN) (Bioanalyzer 6000, Agilent), with an average of 8.5; average RNA yields were 290 ng/µL. The 48 selected samples consisted of four groups: Panic pre-iCBT (n = 12), Panic post-iCBT (n = 11), Control pre (n = 13), and Control post (n = 12) (Supplementary Fig. 1a). Libraries were generated from 500 ng RNA (NEBNext Multiplex Small RNA Library Prep Set for Illumina, New England Biolabs, E7560S) and the small RNA fraction was separated by gel electrophoresis (Invitrogen, G401004) and verified by TapeStation (Agilent Technologies). Libraries were pooled and sequenced on a P4 flow cell (NextSeq 2000 P4 XLEAP-SBS Reagent Kit, 20100995, Illumina) using the NextSeq 2000 system (Illumina) at the Center for Genomic Technologies, the Hebrew University of Jerusalem. Sequence quality control was assessed using FastQC version 0.11.8^49^ and MultiQC version 1.28^50^, with an average final sequencing depth of 6.8M reads per sample. Flexbar program^51^ was then used for adaptor cleaning and additional quality screening. Alignment to miRs was performed with the miRDeep2 algorithm^52^ against miRBase 22.1^53^. Alignment to tRFs was performed using MINTmap^36^ against MINTbase v2.0^37^ and later converted to tDR names using the tDRnamer algorithm^54^.

### qPCR measurements

cDNA synthesis from small RNAs was performed using the RNA Poly(A) Tailing Kit (MCLAB, RPTK-200) and qScript Flex cDNA Synthesis Kit (Quantabio, 95049) with a proprietary oligo(dT) adaptor primer, followed by qPCR using PerfeCTa SYBR Green FastMix Low ROX (Quantabio, 95074), specific forward primers, and a proprietary universal reverse primer. For mRNAs, cDNA synthesis and RT-qPCR were performed using qScript cDNA Synthesis Kit (Quantabio, 95047) and PerfeCTa SYBR Green FastMix (Quantabio, 95072) with human-specific primers. Quantification was performed with the CFX384 Touch Real-Time PCR System (Bio-Rad), and Cq values were extracted with the CFX Maestro software (Bio-Rad v4.1.2433.1219), followed by calculations and plots in R version 4.5.0^55^. Data are presented either as relative expression compared with the housekeeping gene (SNORD47 for small RNAs and b-Actin for mRNA; as Δ*Cq* (*Cq_tar_*_$*et*_ − *Cq*_ℎ*ousekeepin*$_)) or as fold change compared with controls (*FC* = 2^−ΔΔ*Cq*^; ΔΔ*Cq* = Δ*Cq_Panic_* − Δ*Cq_Control_*). In all qPCR plots, outliers were removed using an IQR-based rule within target-, sex-, and study-time-point groups, applied only to groups with at least five observations. Primer sequences are available in Supplementary Table 2.

For tRFs longer than 30nt, a 20-22nt version was designed as a primer. For the Nuc-Val-3’-tRF family, two sequences were chosen as primers. The first was tDR-61-76-Val-AAC-1-M7, since it is the shortest member of the family (16nt long) with 15 of the 16 nts shared among family members (Fig. 5a) and it is one of the ‘transient effect’ tRFs resulting from analysis of individual tRFs (see Fig. 4b). The second, tDR-59-76-Val-AAC-1-M6, was selected as a longer representative of the family (18 nt), based on its shared sequence and high expression (see Fig. 5a,d).

### Statistical analysis

#### Differential expression analysis

As a first quality-control step for the sequencing results, we identified outliers for each small RNA type separately by performing PCA on the raw counts, and excluded samples with a standard deviation larger than 2.5 in at least one PCA for both RNA types. This disqualified five samples in total, resulting in smaller groups, as follows: panic pre-iCBT (n = 10), panic post-iCBT (n = 11), control pre (n = 12), and control post group (n = 10) (Supplementary Fig. 1b, c). The Dream package^56^ in R was used to perform differential expression (DE) analysis for miRs and tRFs separately, and all analyses were performed in R version 4.5.0^55^. First, data were filtered using the filterByExpr function from the edgeR R package^57^, leaving 402 miRs and 677 tRFs in the analysis. Filtered counts were normalized by the Dream package and DE findings were deemed statistically significant if they reached FDR ≤ 0.05. DE analysis was corrected for repeated participant measures, age and BMI (Supplementary Fig. 1d, e), and six comparisons were conducted: (1) panic vs control pre-iCBT, (2) panic post-iCBT vs panic pre-iCBT, (3) panic vs control post-iCBT, (4) control group post- vs pre-equivalent time lapse with no intervention, and two more complex comparisons testing (5) the panic disorder main effect [(panic pre-iCBT + panic post-iCBT)/2 - (control pre + control post)/2] and (6) the iCBT-specific effect [(panic post-iCBT – panic pre-iCBT) - (control post – control pre)].

#### Expression-pattern classification of top-ranked miRs and tRFs

To analyze the expression patterns of top-ranked miRs and tRFs across DE comparisons, we performed the following steps, for both miRs and tRFs, separately. We selected for each comparison the top ten ranked RNAs based on p.value, five of which increased and five decreased based on log2 fold change (log2FC), yielding 43 miRs and 43 tRFs. Next, we excluded RNAs showing a time-related change in controls that was defined as DE analysis log2FC absolute value of 0.5 or higher in the control post-versus pre-time lapse comparison, leaving 32 miRs and 21 tRFs. These remaining top-ranked RNAs were then classified according to the direction and relative magnitude of their log2FC values across the five remaining DE comparisons: (1) panic vs. control at baseline, (2) panic group post- vs. pre-iCBT, (3) panic vs. control post-iCBT, (4) panic disorder main effect and (5) iCBT-specific effect. The log2FC was considered as having an effect if it had an absolute value of 0.4 or above, and as not changing at all if it was 0.15 or lower. RNAs were assigned to a “transient effect” group when the panic-related comparisons (1,4) and the iCBT-related comparisons (2,5) showed opposite directions of change, consistent with a panic-associated alteration that followed iCBT. Specifically, this required concordant directionality between the baseline panic versus control and panic disorder main-effect comparisons, and the opposite concordant directionality between the panic post-versus pre-iCBT and iCBT-specific comparisons.

RNAs were classified as showing an “iCBT effect” when the two iCBT-related comparisons (2, 5) showed concordant changes of at least |log2FC| ≥ 0.4, while the panic-related comparisons (1, 3, 4) showed no change. However, no RNA matched this classification. Conversely, RNAs were classified as showing a “panic effect“ when the baseline panic versus control comparison (1) and the panic disorder main-effect comparison (4) showed concordant changes of at least |log2FC| ≥ 0.4, while the iCBT-related comparisons (2, 5) showed no change. RNAs could also be assigned to the iCBT-effect or panic-effect classifications when the mean absolute effect size in the relevant comparisons exceeded that of the other comparisons by at least 0.25 (the difference between the “change” cutoff of 0.4 and the “no change” cutoff of 0.15). RNAs showing the same direction of change across all examined comparisons were classified as having a “stable effect”. RNAs that did not match any of the previous categories mentioned were retained as “unclassified”. This classification was assigned to RNAs that had an absolute log2FC value ≤ 0.15 across all comparisons, regardless of their initial assignment. The above classifications yielded 13 final miRs and 13 final tRFs for downstream analyses.

#### Multiple regression analysis linking small RNA expression to anxiety severity

Regression analyses were restricted to panic disorder participants. For each miR or tRF target, pre-iCBT, post-iCBT, and combined pre-/post-iCBT analyses were performed using PDSS as the dependent variable. Single time-point associations were determined with baseline, post-session, three-, six-, and 12-month follow-up PDSS scores, as well as PDSS score differences between pre- and post-iCBT PDSS. The combined model used the continuous PDSS composite as the dependent variable that had been additionally adjusted for patients’ time point. Single time-point analyses used ordinary least-squares regression models of the form PDSS ∼ delta Cq + age + BMI + sex. The combined pre-/post-iCBT analysis used a linear mixed-effects model with the same fixed effects and a random intercept for participant identity, PDSS ∼ delta Cq + age + BMI + sex + (1 | participant), allowing participants with only one available time point to remain in the analysis. Models were fitted using complete-case data and were excluded when fewer than 15 participants were available, which led to the exclusion of a male-only analysis. Expression level outliers were removed using an IQR-based rule within target-, sex-, and study-time-point groups, applied only to groups with at least five observations. Benjamini–Hochberg FDR correction was applied across all fitted target-by-time-point tests, and results were deemed significant if FDR ≤ 0.05.

#### Functional enrichment analysis of miR and tRF target genes

Validated target genes were retrieved from miRTarBase^58^ for the 13 top-ranked miRs of the panic effect and transient effect miR groups, comprising five increased miRs (hsa-miR-15a-5p, hsa-miR-132-5p, hsa-miR-132-3p, hsa-miR-23a-5p, hsa-miR-18a-3p) and eight decreased miRs (hsa-miR-543, hsa-miR-125b-5p, hsa-miR-503-5p, hsa-miR-99a-5p, hsa-miR-331-5p, hsa-let-7e-5p, hsa-miR-99b-3p, hsa-miR-412-5p), based on the panic pre-iCBT versus control pre comparison. Functional enrichment analysis was performed in R using the clusterProfiler package^59^, with each miR target set analyzed separately to resolve miR-specific functional signatures. Enrichment testing was carried out using WikiPathways with the enrichWP package^38^. Multiple testing correction was performed using the Benjamini-Hochberg procedure, with an adjusted p-value threshold of 0.05. For tRFs, tRFtar^39^ was used to identify the validated targets of the 11 members of the Nuc-Val-3’-tRF family, and subsequent WikiPathway analysis was done within the tRFtar website. The results were later combined with the miR results for downstream analysis.

To elucidate the potential regulatory effect on immune-related processes of PBMC small RNAs in panic disorder and iCBT response, we constructed a bipartite network linking together (i) the panic- and transient-effect miRs and the Nuc-Val-3’-tRF family members with (ii) their enriched biological pathways. We annotated the significant WikiPathways found previously to be immune- or inflammation-related, which accounted for 15.3% of the identified pathways. The bipartite network was generated using a force-directed stress layout implemented in the R package ggraph (v2.1.0)^60^.

### Use of large language models

ChatGPT (GPT-5.6 Sol, OpenAI; accessed August 2026) was used for language refinement during manuscript preparation. Google Antigravity (Gemini 3.1 Pro, Google; accessed April 2026) was used to assist with code refinement and debugging. AI-assisted outputs were reviewed and validated by the authors, who take full responsibility for the content of the manuscript and the analyses.

