## Supplementary Information for "Internet-delivered cognitive behavioral therapy is associated with distinct blood small RNA profiles in panic disorder"

### Supplementary Results

#### Responsive miRs distinguish panic disorder and post-iCBT states in PCA

PCA based on the 13 responsive miRs separated panic disorder patients from controls and distinguished the pre- and post-iCBT panic groups (Fig. 2c). The only cross-group overlap involved paired pre- and post-iCBT samples from the same participant. PC loading analysis identified hsa-miR-412-5p as the main contributor to patient-control separation on PC1 (Supplementary Fig. 3a), whereas hsa-miR-23a-5p contributed most to the pre- versus post-iCBT separation on PC2 (Supplementary Fig. 3b). These findings indicate that the selected miR set captured both panic disorder-associated differences and a distinct post-treatment molecular state.

#### miR profiles differ between panic participants who did and did not achieve remission

To assess selected miRs in broader clinical subgroups, we quantified candidate miRs by qPCR in the full cohort, including male and female patients and post-iCBT remitters and non-remitters. Because all male patients achieved remission, non-responders could be evaluated only among females. Candidate miRs were selected from the strongest PC contributors and from the panic- and transient-effect groups with a sequencing base mean expression above five CPM (Fig. 2b, c; Supplementary Fig. 3a, b), yielding hsa-let-7e-5p, hsa-miR-125b-5p, hsa-miR-15a-5p and hsa-miR-23a-5p (Supplementary Fig. 3c). In female remitters, all four miRs followed the same direction of change observed in the sequencing cohort. Group differences were generally modest; however, female remitters and non-remitters showed significant miR-specific differences for hsa-miR-125b-5p (Supplementary Fig. 3c).

#### Top miR target genes highlight immune- and inflammation-related pathways

Target enrichment analysis of the 13 panic- and transient-effect miRs (Fig. 2b) showed that ~15% of significantly enriched pathways were immune-related (Supplementary Fig. 8), consistent with inflammatory alterations reported in panic disorder[1,2]. The pathway shared by the largest number of miRs was miR-targeted genes in lymphocytes, involving hsa-let-7e-5p, hsa-miR-125b-5p, hsa-miR-412-5p, hsa-miR-503-5p, hsa-miR-543 and hsa-miR-15a-5p. Additional enriched pathways included TNF-alpha, TGF-beta, JAK-STAT and MAPK signaling. BDNF signaling was also detected for hsa-miR-125b-5p, hsa-miR-132-3p and hsa-miR-15a-5p, in line with evidence for altered blood BDNF in panic disorder[3]; hsa-miR-125b-5p and hsa-miR-132-3p also target acetylcholinesterase, among other transcripts[4,5]. Finally, FSH- and estrogen-related pathways were identified among selected miR targets, consistent with reported effects of ovarian hormone signaling on anxiety-related behaviour[6]. Overall, these enrichments support an immune- and hormone-related functional context for miRs that distinguish panic disorder and iCBT-response states.

#### Increased and decreased small RNAs co-target immune-regulatory pathways

Among small RNAs increased in panic patients before iCBT (Fig 2c, Supplementary Fig. 8), hsa-miR-15a-5p was the largest network hub, with 12 pathway connections spanning lymphocyte and receptor-signaling processes, including B-cell receptor, T-cell receptor, IL-1 and IL-18 signaling. hsa-miR-132-5p targeted pathways centered on innate immune activation and neuroinflammation and was the only individual small RNA to show significant global immune-pathway enrichment (Fisher test, 80%, BH-adjusted $p = 0.006$). The Nuc-Val-3'-tRF family also emerged as a major hub, linked to interleukin, inflammatory-response and type I interferon pathways; more than half of these pathways were shared with one to four miRs.

Conversely, decreased miRs, including hsa-miR-125b-5p and hsa-miR-543, were linked to overlapping chronic inflammation, cytokine, JAK-STAT, TNF-alpha, neuroinflammation and chemokine-related pathways. Several pathways were co-targeted by both increased and decreased small RNAs, including lymphocyte, IL-5, TNF-alpha, IL-4 and macrophage-stimulating protein signaling. This overlapping target structure suggests that panic disorder- and iCBT-responsive small RNAs may converge on shared immune pathways while exerting complementary regulation of systemic inflammation, lymphocyte activation and cytokine signaling in PBMCs.

### Supplementary Figures

#### Supplementary Figure 1

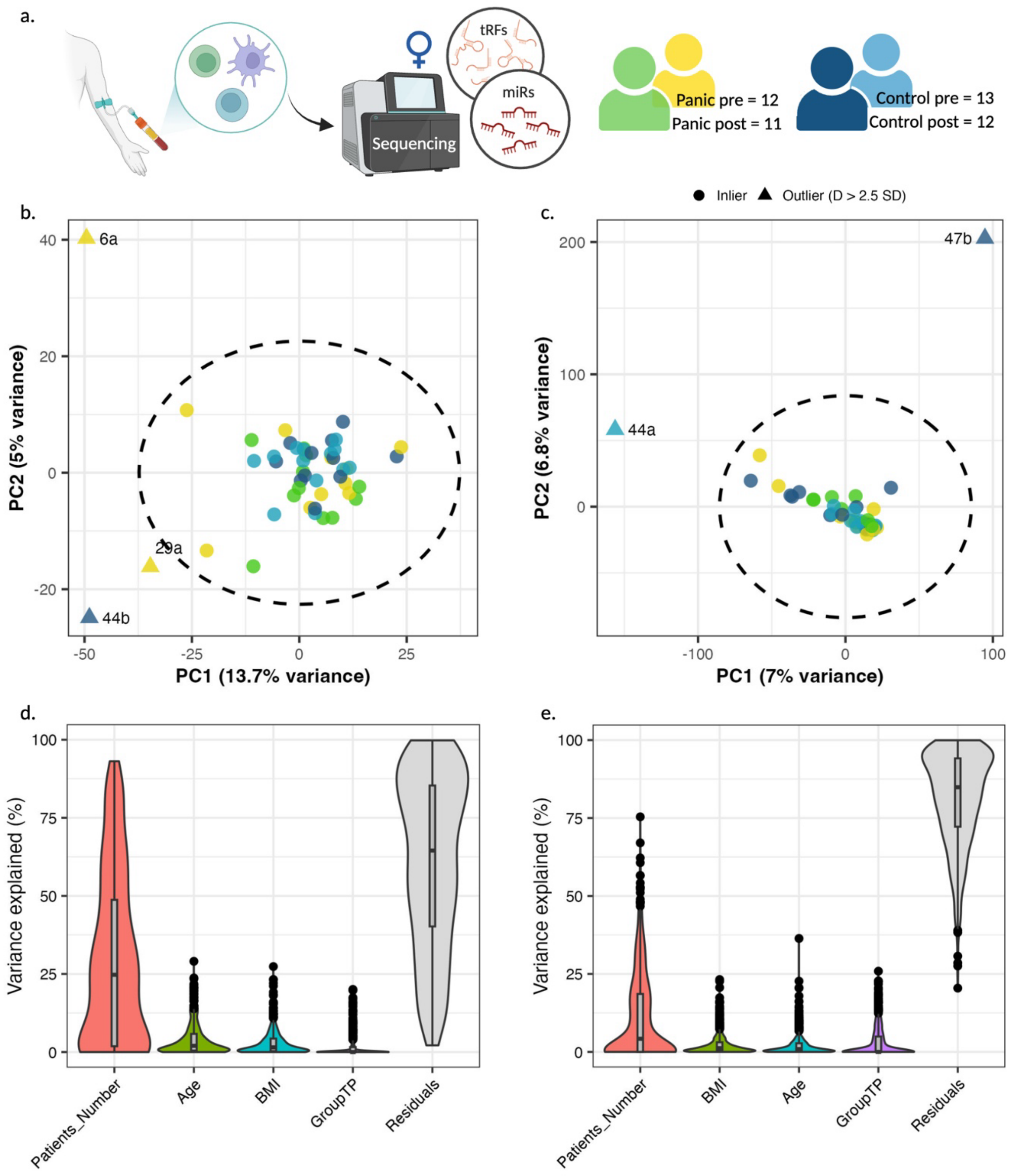

**Outlier selection and variance partitioning in the miR and tRF sequencing cohorts.** (a) Blood was drawn from participants at two time points (see Fig.1a), RNA was extracted, and only female panic patients who achieved remission at the post-iCBT session and their matched controls were sequenced (see Methods section). (b,c) PCA plot of raw counts per participant for (b) miRs and (c) for tRFs. In both cases outliers were identified using a 2.5 standard deviation (SD) cutoff. (d,e) Violin plot showing the fraction of variance explained by each variable in the design model for (d) miRs and (e) for tRFs. In both cases each point represents one sample. Created with Biorender.

#### Supplementary Figure 2

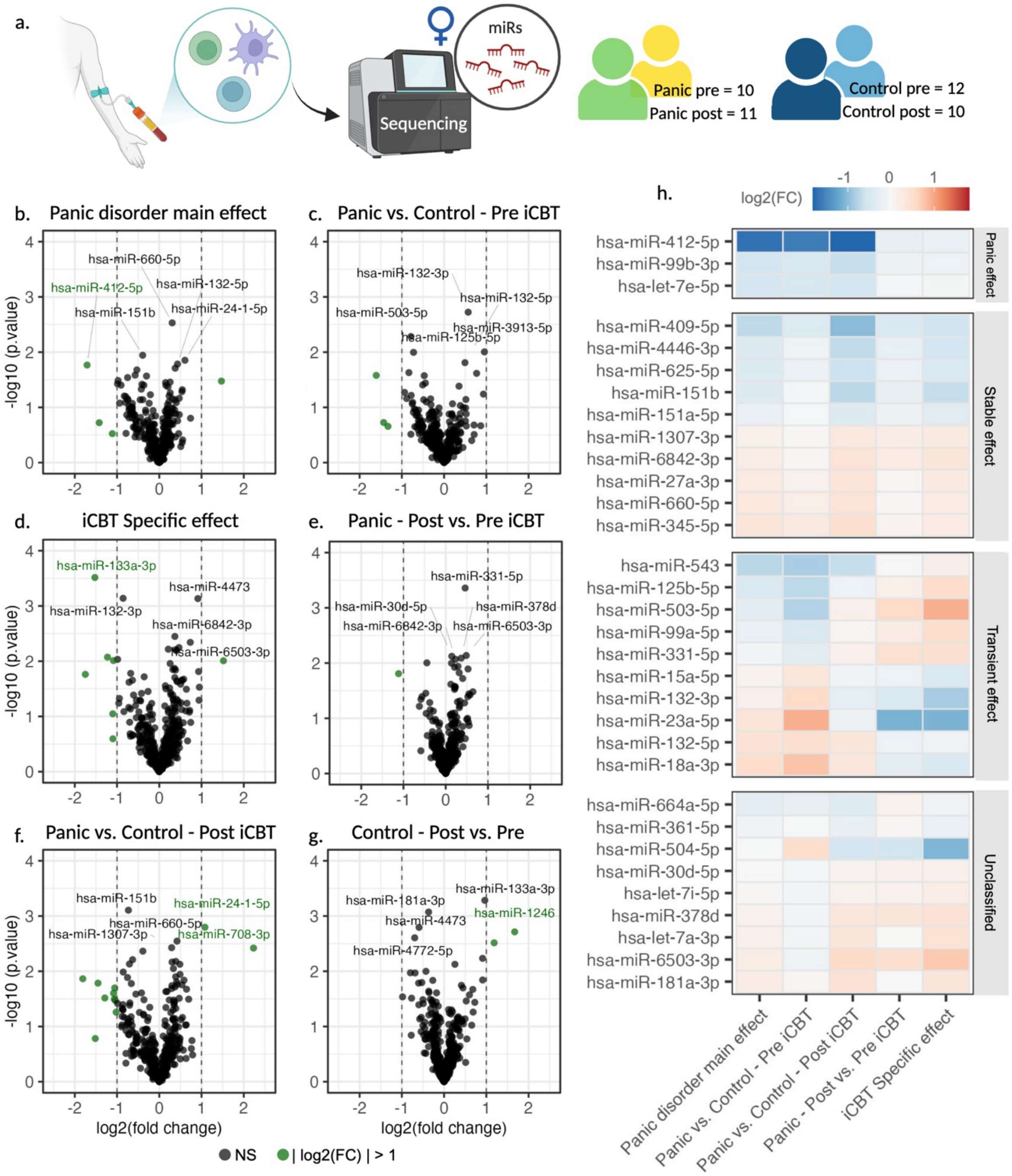

**Expression-pattern grouping distinguishes miR effects across comparisons. (**a) RNA was extracted from PBMCs and sequenced; analyses shown include only female panic patients who achieved remission after iCBT and their matched controls. (b–g) Volcano plots showing miR differential-expression results for six comparisons: (b) Panic main effect, (c) Panic vs. control pre-iCBT, (d) iCBT-specific effect, (e) Panic post- vs. pre-iCBT, (f) Panic vs. control post-iCBT, and (g) controls post- vs. pre-similar interval without intervention. Black labels indicate non-significant miRs, with log2(FC) <1, and green labels indicate miRs with log2(FC) >1. Named labels indicates the four top ranked miRs in term of p.value in each comparison. No miR reached significant level after FDR. (h) Heatmap of the top 10 ranked miRs in each of the five indicated comparisons, excluding the control only comparison. miRs with |log2FC| ≥ 0.5 in the control post- vs. pre-comparison were excluded and the remaining miRs were grouped into four categories determined by differential expression patterns across the five comparisons (Panic effect, also shown in Figure 2; Stable effect; Transient effect, also shown in Figure 2; Unclassified). Color indicates log2 fold change (log2FC) increase (red) or decrease (blue) for each miR in the indicated comparison. Created with Biorender.

#### Supplementary Figure 3

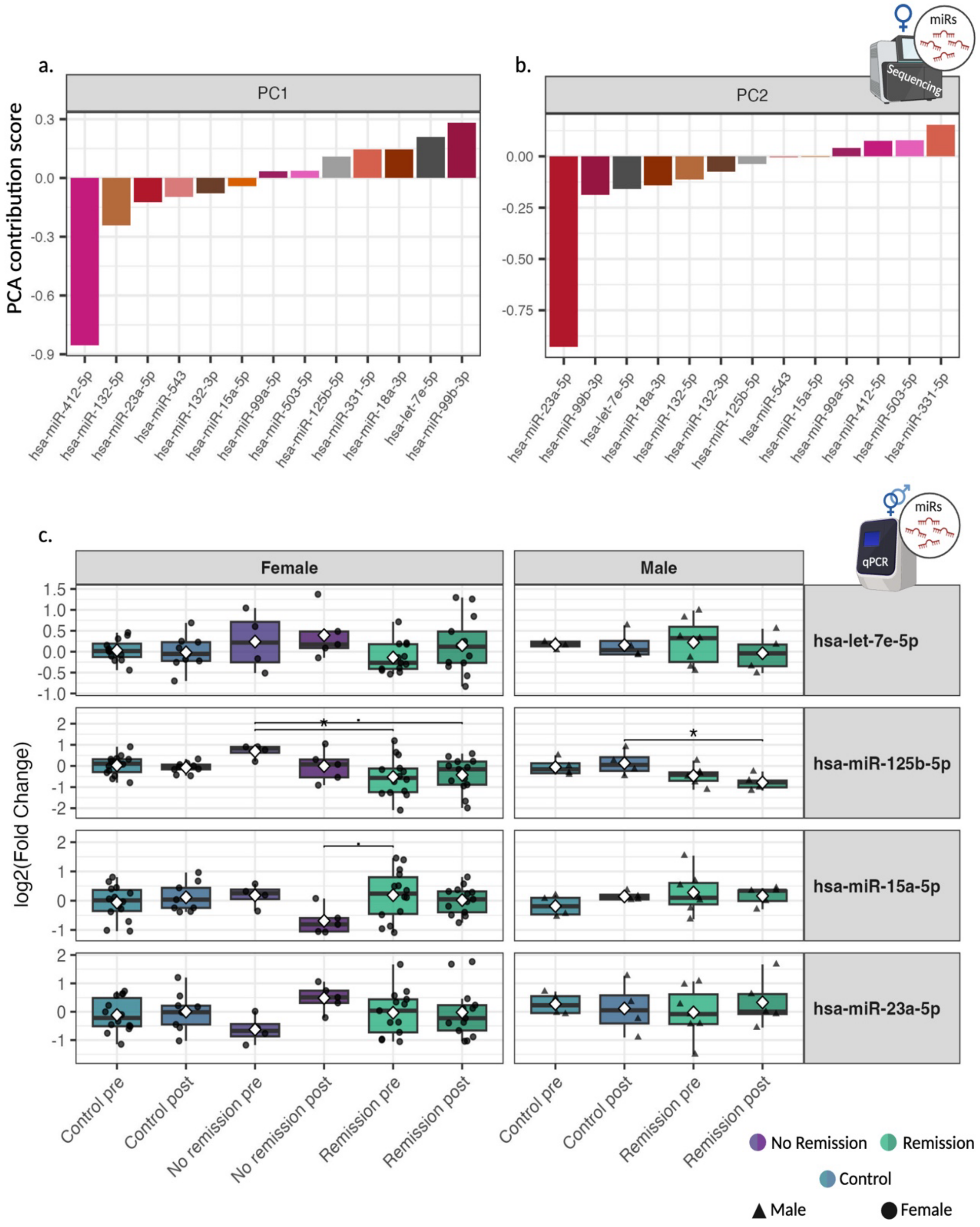

**Contributions of individual miR to PCA structure and their levels in the full cohort.** (a,b) Bar plots showing the contribution score of each of the 13 miRs to the construction of the two PCs shown in Fig. 2, (a) PC1 and (b) PC2. Each mir is colored in a different color bar. Sample sizes: panic pre-iCBT n = 10, panic post-iCBT n = 11, control pre n = 12, control post n = 10. (c) Boxplots showing levels of four miRs in female and male participants of the full cohort, determined by qPCR. Results are shown as log2FC relative to controls across sex, group and post-session remission status. Statistics represent Tukey–Kramer tests following ANOVA, with FDR ≤ 0.05 (. = p≤0.1, * = p≤0.05). White diamond in each box indicates mean and the horizontal line marks the median. qPCR was performed only for miRs with CPM > 5 in the sequencing results, selected from miRs with the highest contribution scores on the two PCs or from the panic- and transient-effect miR groups in Fig. 2. Sample sizes by group for pre and post time points, respectively: females, remission n = 14/13, no remission n = 4/5, controls n = 14/10; males, remission n = 7/5, no remission n = 0/0, controls n = 4/4. Created with Biorender.

#### Supplementary Figure 4

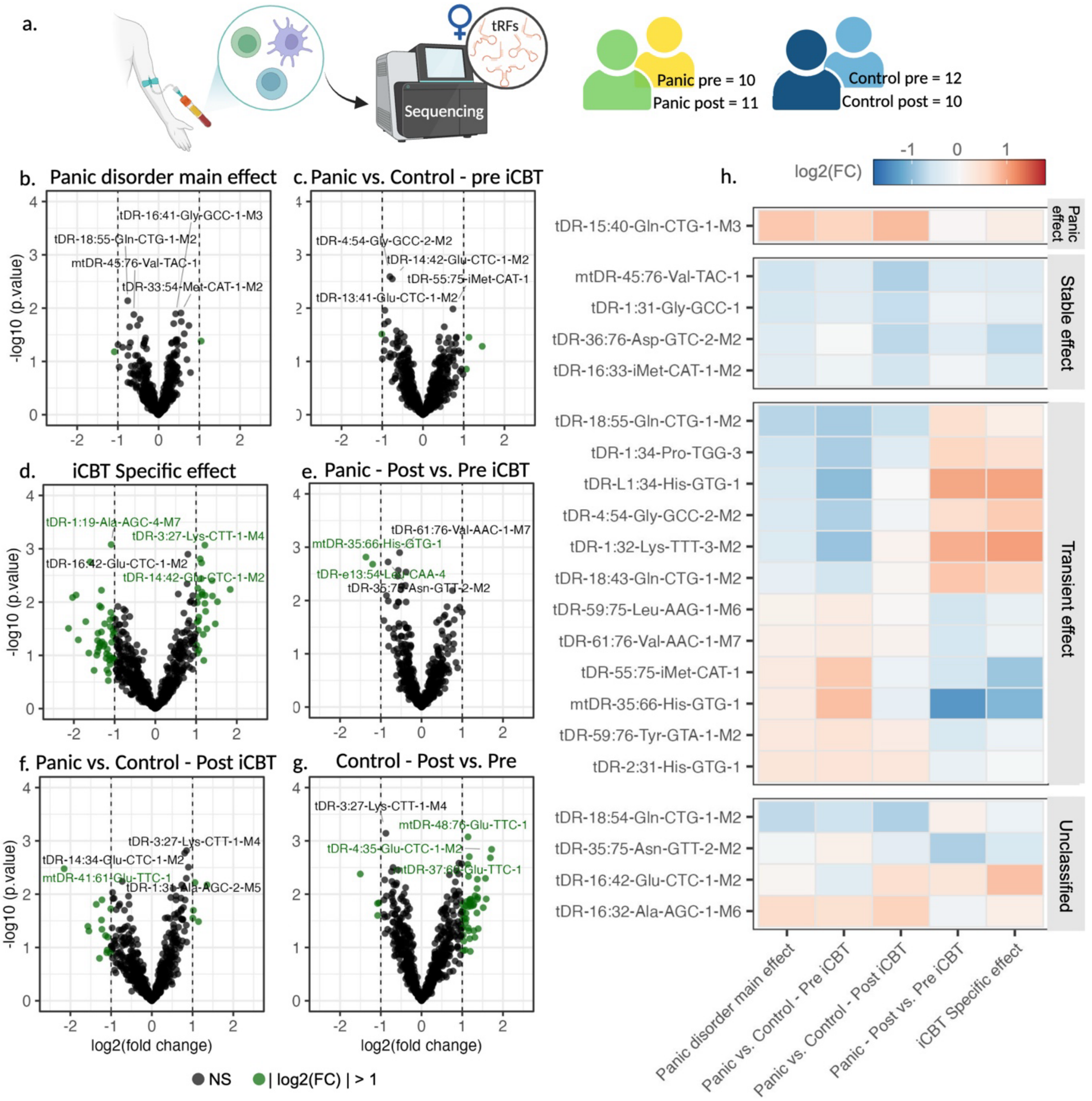

**Expression-pattern grouping distinguishes tRF effects across comparisons**. (a) RNA was extracted from PBMCs and sequenced; analyses shown include only female panic patients who achieved remission after iCBT and their matched controls. (b–g) Volcano plots showing tRF differential expression results for six comparisons: (b) Panic main effect, (c) Panic vs. control at baseline, (d) iCBT-specific effect, (e) Panic post- vs. pre-iCBT, (f) Panic vs. control post-iCBT, and (g) controls post- vs. pre-similar interval without intervention. Black labels indicate non-significant tRFs, with log2(FC) <1, and green labels indicate tRFs with log2(FC) >1. Named labels indicates the four top ranked tRFs in term of p.value in each comparison. No tRF reached significant level after FDR. (h) Heatmap of the top 10 ranked tRFs from each of the five indicated comparisons. Color indicates log2 fold change (log2FC) increase (red) or decrease (blue) for each tRF in the indicated comparison. tRFs with |log2FC| ≥ 0.5 in the control post- vs. pre-comparison were excluded, and the remaining tRFs were grouped by differential-expression pattern across the five comparisons. Created with Biorender.

#### Supplementary Figure 5

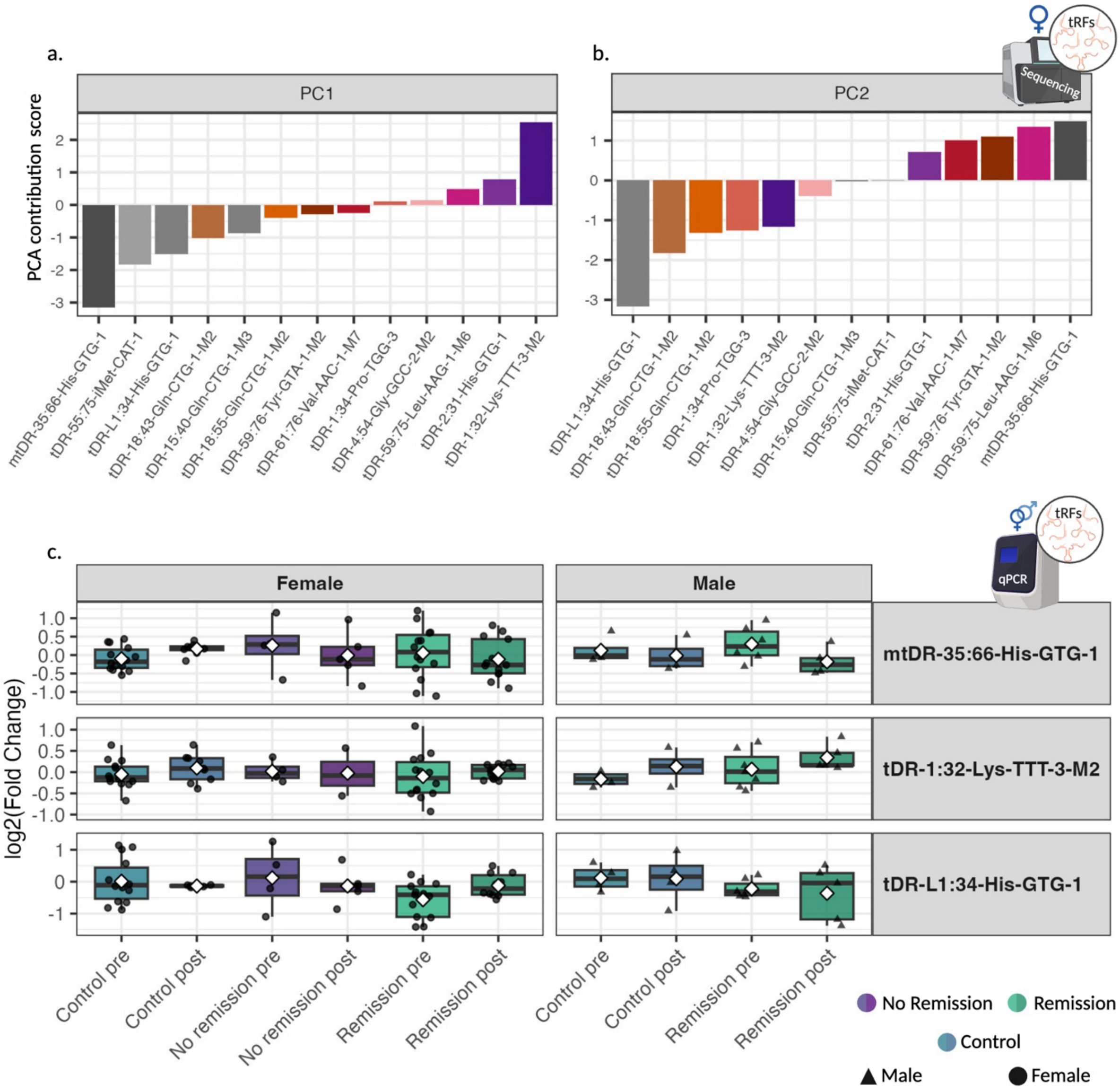

**Individual tRF contributions to PCA structure, and levels in the full cohort**. (a,b) Bar plots showing the contribution score of each tRF to the construction of the two PCs shown in Fig. 4: (a) PC1 and (b) PC2. Sample sizes: panic pre-iCBT n = 10, panic post-iCBT n = 11, control pre n = 12, control post n = 10. (c ) Boxplots showing tRF log2FC relative to controls across sex, group and post-session remission status, based on qPCR results from male and female participants in the full cohort. Statistics represent Tukey–Kramer tests following ANOVA, with FDR ≤ 0.05. tRF candidates for qPCR were selected from tRFs with the highest contribution scores on the two PCs. Sample sizes by group for pre/post time points, respectively: females, remission n = 14/13, no remission n = 4/5, controls n = 14/10; males, remission n = 7/5, no remission n = 0/0, controls n = 4/4. Created with Biorender.

#### Supplementary Fig. 6

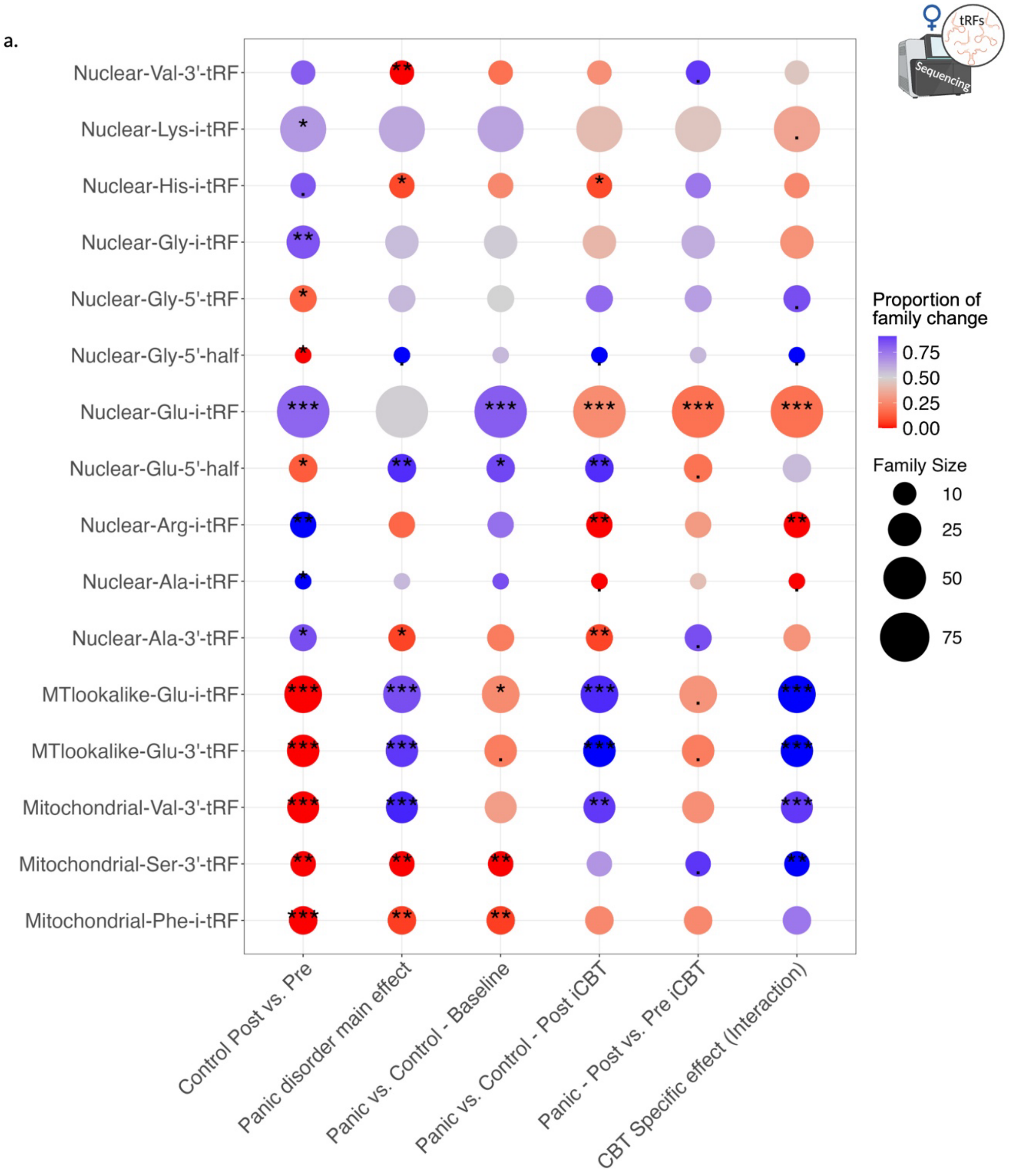

**tRF families differentiate between study groups, with high variability.** (a) Dot plot of tRF families showing directional family-level changes assessed with exact binomial tests. Dot size indicates the number of individual tRFs within each family and color indicates the proportion of family members increased or decreased in each differential expression comparison. Only families with FDR ≤ 0.05 in at least one of the six comparisons are shown (. = p≤0.1, * = p≤0.05, ** = p≤0.01, *** = p≤0.001). Sample sizes: panic pre-iCBT n = 10, panic post-iCBT n = 11, control pre n = 12, control post n = 10. Created with Biorender.

#### Supplementary Fig. 7

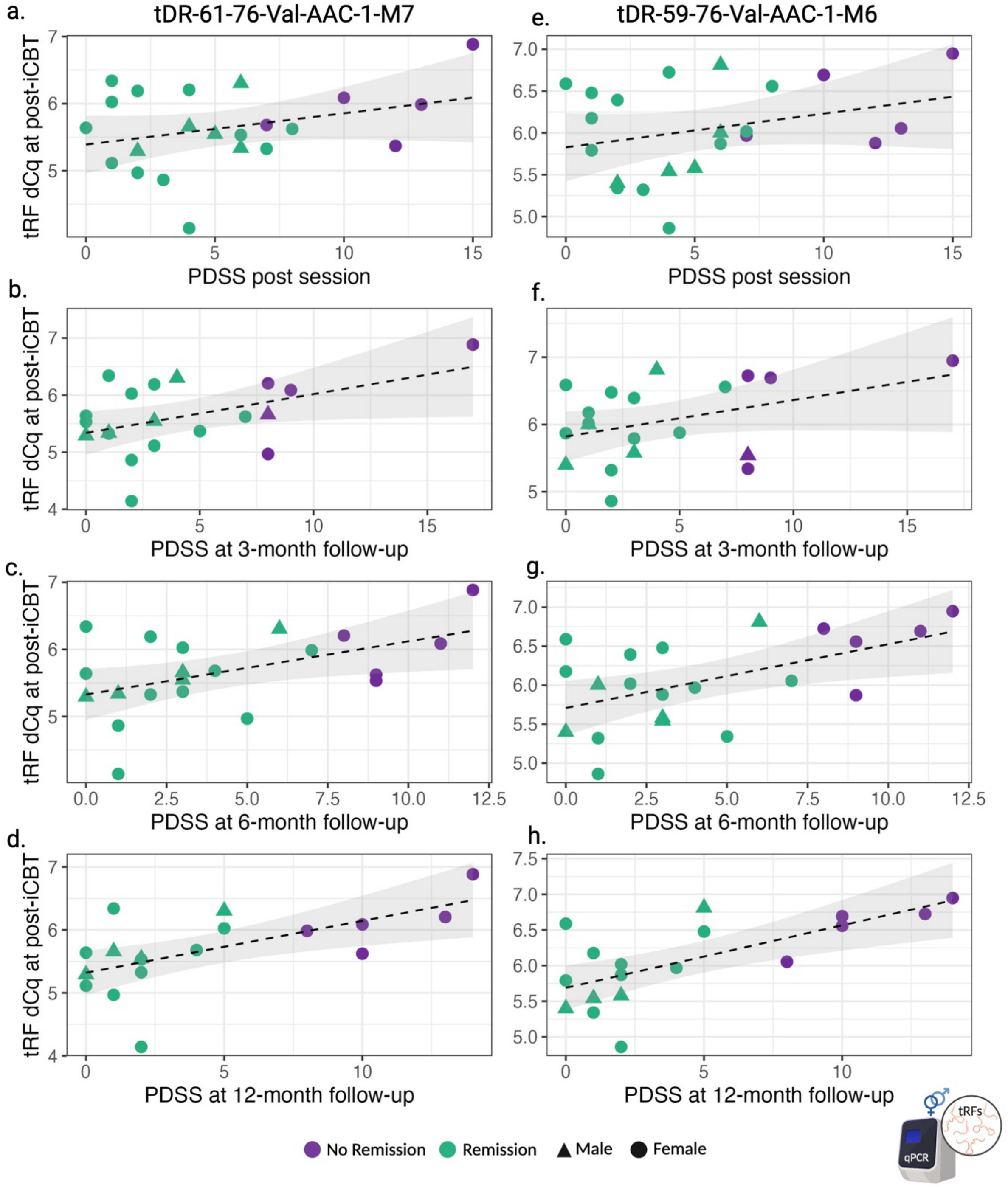

**Post-iCBT Nuclear-Val-3′-tRF levels are associated with PDSS at follow-up interviews.** Scatter plots showing post-treatment ΔCq values for two members of the Nuclear-Val-3′-tRF family, tDR-61-76-Val-AAC-1-M7 (a–d) and tDR-59-76-Val-AAC-1-M6 (e–h), in relation to PDSS at post-iCBT session (a,e), three-month follow-up interview (b,f), six-month follow-up interview (c,g) and 12-month follow-up interview (d,h). Points are colored by remission status at the corresponding PDSS evaluation, with females in circles and males in triangles. Associations were tested using multiple linear regression models with age, sex and BMI as covariates. Model fit is reported as Δadjusted $R^2$ and FDR: (a) Δadjusted $R^2$ = 0.062, FDR = 0.214; (b) Δadjusted $R^2$ = 0.21, FDR = 0.066; (c) Δadjusted $R^2$ = 0.21, FDR = 0.066; (d) Δadjusted $R^2$ = 0.42, FDR = 0.038; (e) Δadjusted $R^2$ = 0.043, FDR = 0.260; (f) Δadjusted $R^2$ = 0.14, FDR = 0.138; (g) Δadjusted $R^2$ = 0.27, FDR =

0.030; (h) Δadjusted $R^2 = 0.44$, FDR = 0.030. Dashed lines indicate fitted regression lines and shaded areas indicate 95% confidence intervals. Sample sizes by group for pre and post time points, respectively: females, remission n = 14/13, no remission n = 4/5, controls n = 14/10; males, remission n = 7/5, no remission n = 0/0, controls n = 4/4. Created with Biorender.

#### Supplementary Fig. 8

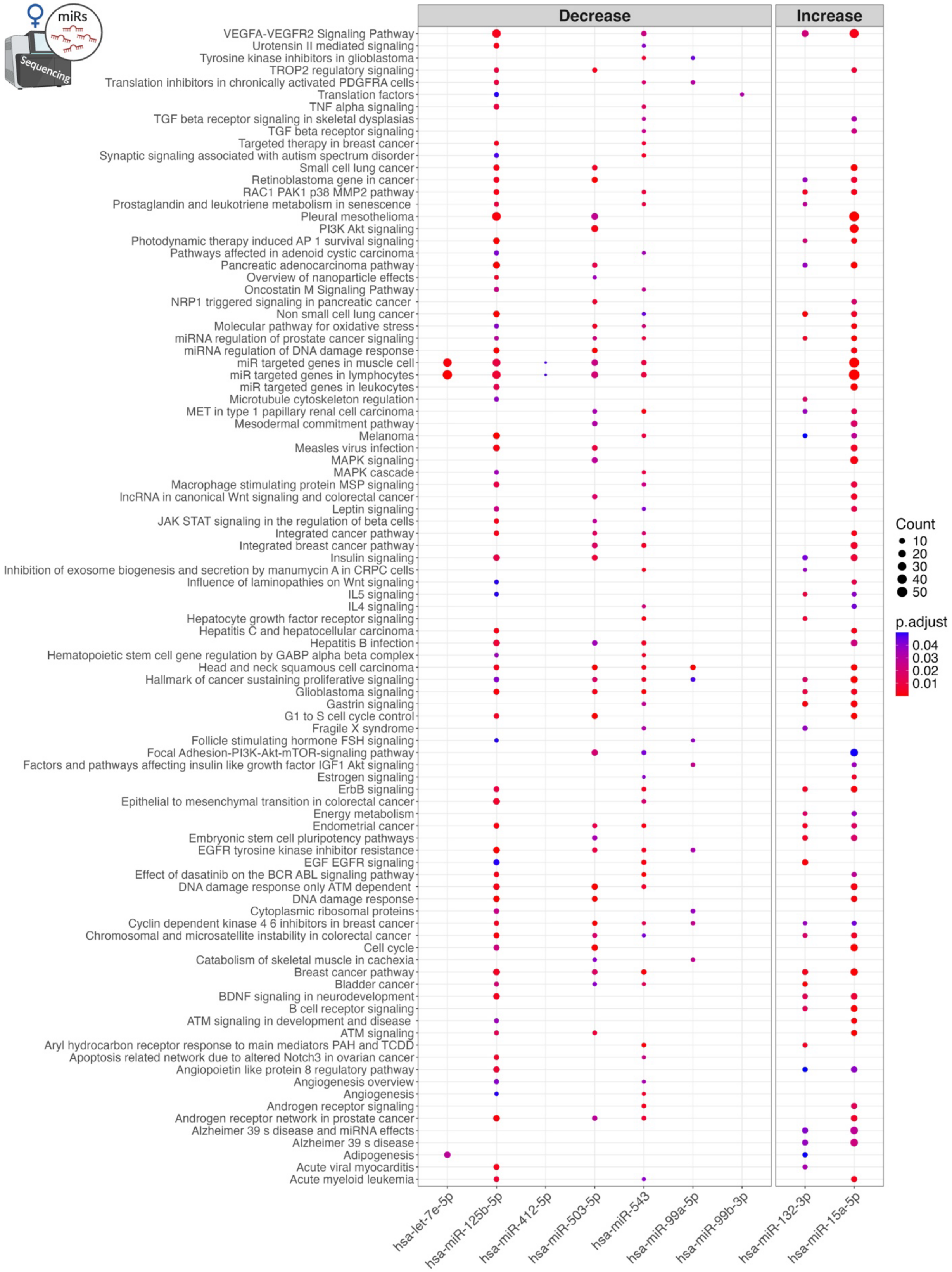

**Top ranking miRs converge on immune-related pathways.** Dot plot showing WikiPathways terms (y-axis) enriched among predicted targets of two or more miRs from the panic-effect and transient-effect groups, restricted to miRs with significant pathway-enrichment results (x-axis), which increase (right) or decrease (left). Dot size indicates the number of target genes contributed by each miR within a given pathway and color indicates the enrichment-test adjusted p-value calculated using the Benjamini–Hochberg correction. Only experimentally validated miR targets from miRTarBase were analyzed. Created with Biorender.

#### Supplementary Fig. 9

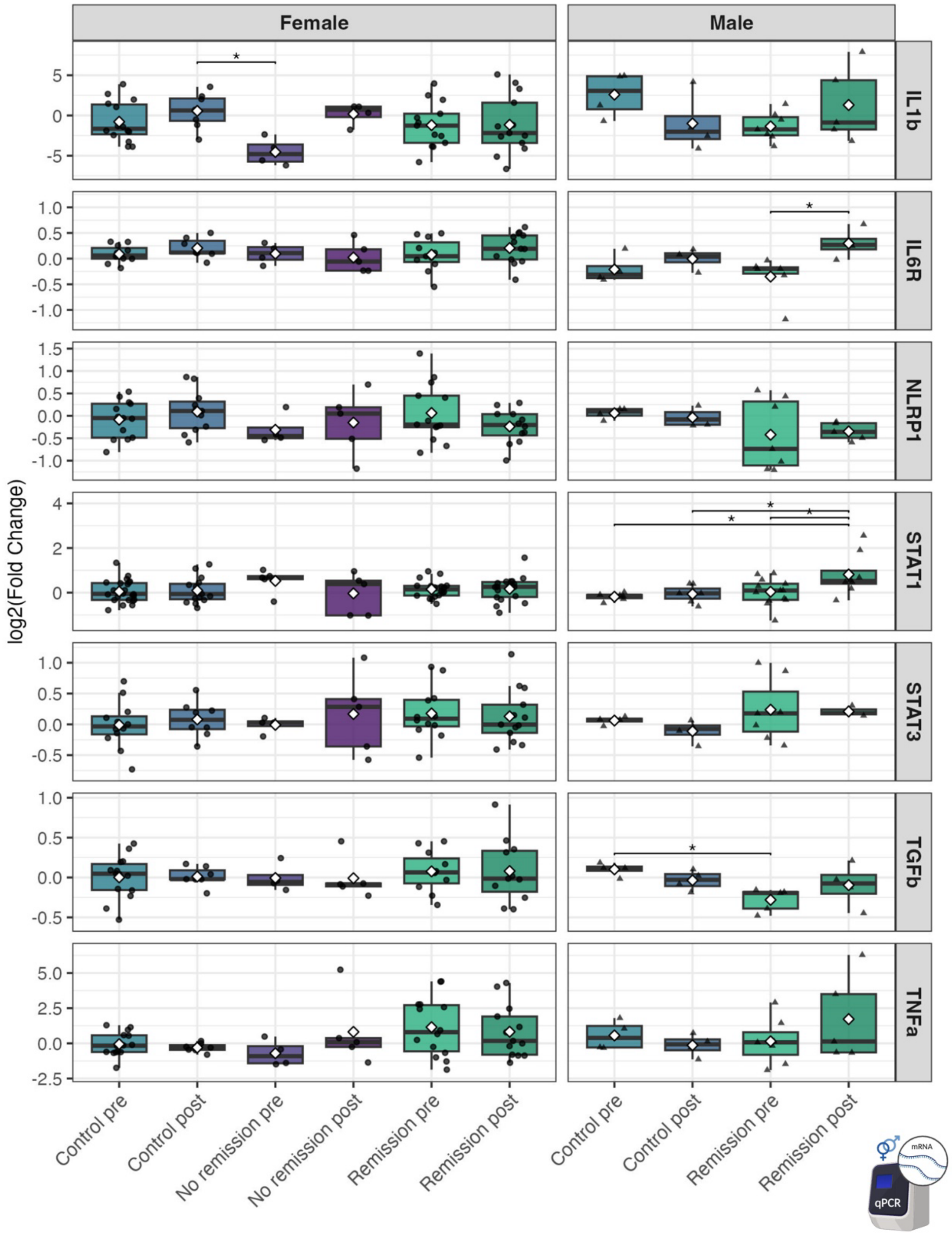

**Immune- and Inflammation-related mRNA markers in the full cohort.** Boxplots showing qPCR results for immune- and inflammation-related mRNA transcripts, using PBMC RNA from the full cohort. Results are presented as log2(fold change) relative to controls across sex, group and post-session remission status. Statistics represent Tukey–Kramer tests following ANOVA, with FDR ≤ 0.05 (. = p≤0.1, * = p≤0.05). White diamond in each box indicates mean and the horizontal line marks the median. Sample sizes by group for pre and post time points, respectively: females, remission n = 14/13, no remission n = 4/5, controls n = 14/10; males, remission n = 7/5, no remission n = 0/0, controls n = 4/4. Created with Biorender.
